# CNVSIM: an interpretable cross-study copy-number reference atlas for cancer subtyping

**DOI:** 10.64898/2026.10.26.761332

**Authors:** Jiahui Yu, Juliette Bardet-Trouilloud, Stephanie Jaeger, Michael Baudis

## Abstract

Genome-wide copy-number variation (CNV) profiles provide a representation of structural changes in cancer genomes, but interpreting new profiles with respect to their underlying biology remains challenging. We developed CNVSIM, which organizes CNV profiles into root-specific reference spaces and defines subtypes with chromosome-scale and gene-level signatures. Applied to the Progenetix dataset, CNVSIM established 112 CNV-defined subtypes across 19 cancer roots, supported by 155,881 neoplastic analyses from diverse studies and assay platforms. In the context of diagnostic concepts, these subtypes distinguish genomic architectures and reveal associated differences in somatic mutation patterns, with selected associations supported across cohorts. In practice, CNVSIM enables new profiles to be projected onto the established reference and linked to corresponding reference analyses. Additionally, CNVSIM enables the identification of cancer cell lines with copy-number architectures similar to individual subtypes, while also revealing gaps in subtype and alteration coverage based on available data. Reference robustness was supported by subtype recovery after data reduction and assignment of 77.6% of held-out TCGA profiles using a reference reconstructed without TCGA.

## INTRODUCTION

Copy-number alterations are a major component of cancer genome variation, ranging from whole-chromosome and chromosome-arm changes to focal amplifications and deletions^1,2^. These alterations are commonly summarized as genome-wide copy-number variation (CNV) profiles for individual samples. Recurrent CNV architectures vary across and within cancer types, making these profiles a useful basis for comparing cancer genomes^3^.

Large public cancer genomics resources, including The Cancer Genome Atlas (TCGA) and cBioPortal, have made CNV profiles increasingly accessible^4,5^. Progenetix integrates CNV data from these and numerous additional public studies into the largest publicly accessible collection of curated, harmonized genome-wide CNV profiles from neoplastic samples^6–8^. Such cross-study integration provides greater reference depth for recurrent CNV architectures that may be sparsely represented in individual cohorts. However, access to a large collection of CNV profiles is not equivalent to having a reusable structural reference. While data repositories enable retrieval by diagnostic and genomic annotations, a reference atlas organizes profiles into recurring CNV-defined groups against which new samples can be compared. Such a reference would support consistent comparisons across studies and provide a framework for examining CNV heterogeneity within established diagnoses.

Reusable reference frameworks have become an effective strategy for organizing complex biological data. Curated cancer knowledge bases such as OncoKB and the Cancer Gene Census from the Catalogue of Somatic Mutations in Cancer (COSMIC) provide shared annotations for cancer genes and their alterations^9,10^. In contrast, single-cell atlases define reference cellular states onto which new query datasets can subsequently be mapped^11,12^. Although serving different purposes, these resources illustrate how shared references can support consistent interpretation and reuse of biological information across independent studies.

Existing CNV studies have addressed different aspects of CNV biology in cancer. Pan-cancer analyses have identified recurrent CNV regions and their frequencies across cancer types^3,13^, whereas copy-number signatures have related genome-wide CNV patterns to chromosomal-instability processes such as aneuploidy, whole-genome doubling, homologous recombination deficiency and chromothripsis^14^. Other studies have organized CNV profiles by classifying them into predefined diagnostic categories^15,16^ or identifying molecular subgroups within individual cancer types or multi-omic cohorts^17,18^. Together, these approaches have established the biological relevance of CNV landscapes, process signatures and molecular subgroups. However, a reusable cross-study reference spanning multiple cancer types is still needed to organize profiles by recurrent CNV architecture and support comparison with new profiles from neoplastic samples and experimental models.

Constructing such a reference from heterogeneous public CNV data presents several challenges. Progenetix integrates CNV profiles generated using hundreds of assay platforms spanning chromosomal comparative genomic hybridization (CGH), array-based and sequencing-based technologies, together with diverse study cohorts and diagnostic annotations^7^. Differences in assay resolution and study composition can introduce technical variation that complicates the identification of recurrent biological structure. Furthermore, diagnostic labels do not map one-to-one onto CNV architecture: a broad diagnosis may encompass multiple recurrent CNV patterns, while similar CNV architectures may occur across different morphological diagnoses^19^. Addressing these challenges requires a reference that captures recurrent biological structure while minimizing technical variation and organizing profile comparisons within a common genomic framework that complements diagnostic classifications.

Here, we present CNVSIM, an interpretable cross-study reference atlas of cancer CNV architecture. Built from Progenetix, the atlas comprises 112 CNV-defined subtypes across 19 anatomical groups, supported by 155,881 defining analyses from multiple assay platforms and studies. These subtypes capture distinct CNV architectures within shared diagnoses and are characterized by chromosome-scale and gene-level signatures. Some also show within-diagnosis somatic mutation associations, although mutation data were not used to define the subtypes. New CNV profiles can be projected onto the atlas to identify a reference subtype and retrieve similar reference analyses together with associated structural and molecular annotations. In held-out evaluations, projected profiles showed high assignment coverage and strong correspondence with the reference subtype structure. Applying the atlas to cancer cell lines identified recommended models for 90 subtypes while revealing characteristic gene-level CNV alterations not recorded in these models. Together, these results establish CNVSIM as a reusable structural reference for cross-study CNV comparison, interpretation of new profiles, and informed selection of candidate experimental models.

## RESULTS

### CNVSIM establishes a cross-study reference of cancer CNV subtypes

CNVSIM was built using CNV profiles from Progenetix, an open cancer genomics resource integrating public CNV studies with harmonized sample-level metadata. Of the 257,365 available analyses, 204,587 analyses remained for CNVSIM evaluation after excluding non-neoplastic reference analyses and analyses with uninformative platform annotations. Cancer cell-line analyses were retained separately for downstream projection. The 204,587 analyses used for atlas construction were grouped into 22 topography (ICD-O-T) roots of the International Classification of Diseases for Oncology, third edition (ICD-O-3)^20^, of which 19 yielded reference CNV subtypes.

The source CNV profiles were generated using 235 assay platforms spanning chromosomal comparative genomic hybridization (CGH), genomic arrays, including single-nucleotide polymorphism (SNP) arrays, and next-generation sequencing (NGS). These technologies detect genomic copy-number imbalances but differ in genomic coverage, resolution and sensitivity to small or low-amplitude alterations^21–24^. For atlas construction, platform annotations were harmonized into three families: NGS-based assays, high-resolution arrays, and low-resolution arrays or chromosomal CGH. Arrays with at least 180,000 probes were classified as high-resolution, whereas lower-density arrays and chromosomal CGH profiles were assigned to the low-resolution family.

To reduce dimensionality while retaining disease-relevant signals, analyses were represented by CNV states across 1,263 autosomal cancer-associated genes curated from OncoKB^9^ and the COSMIC Cancer Gene Census^10^.

The assembled dataset combined different assay platforms and varied substantially across cancer roots in analysis count, CNV burden, genome-wide alteration fraction and cross-platform agreement of gene-level CNV frequencies (Figure 1A). These observations motivated a rootwise rather than platform-stratified design. A single pan-cancer comparison space would combine samples from anatomical sites with distinct CNV architectures, whereas pre-stratifying by platform, source or fine-grained diagnostic labels would fragment the data and reduce the ability to identify recurrent CNV patterns shared across platforms. CNVSIM therefore constructed root-specific comparison spaces within grouped ICD-O-T roots, using platform associations to evaluate candidate graph configurations while retaining assay-platform and data-source composition as annotations for each reference subtype.

**Figure 1.**
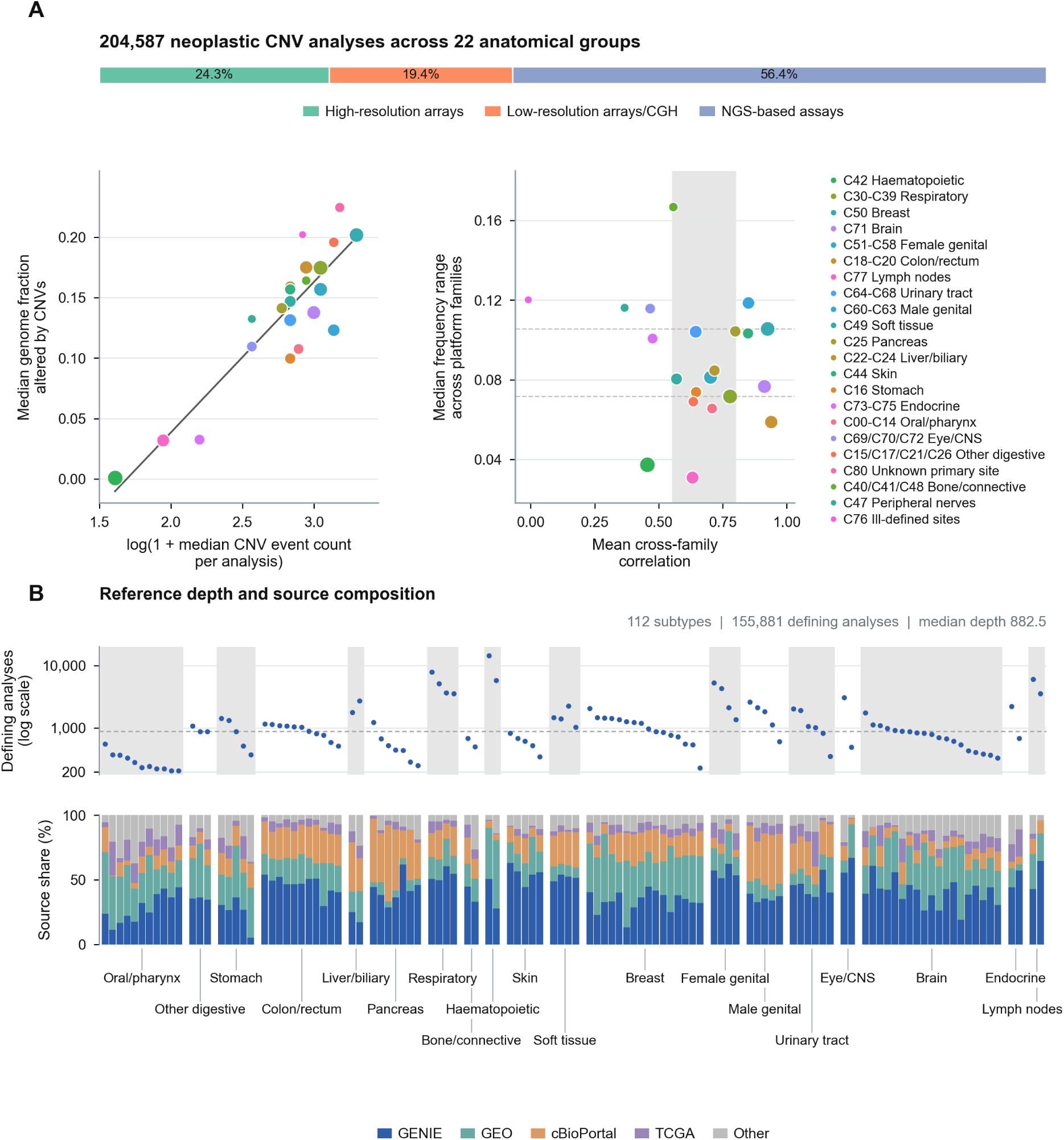
Heterogeneity of CNV profiles across cancer roots and composition of the CNVSIM reference atlas. **(A)** Assay-platform family distribution, root-wise CNV characteristics and cross-platform consistency across the 204,587 neoplastic analyses from 22 cancer roots. The top bar shows the overall proportions of high-resolution arrays, low-resolution array/CGH profiles and NGS-based profiles. Below, each point represents one cancer root, with point size proportional to the number of analyses. Left, the median CNV event count per analysis (log(1+count)) is plotted against the median genome fraction altered by CNVs; the line indicates the fitted cross-root linear trend. Right, the mean pairwise Pearson correlation of autosomal gene-level CNV frequencies between platform families is plotted against the median across-gene difference between the maximum and minimum platform-family frequencies. The vertical grey band and horizontal dashed lines indicate the interquartile ranges of cross-root correlation and frequency divergence. **(B)** Reference depth and data-source composition across the 112 reference subtypes. In the upper track, each point represents one subtype, showing the number of defining analyses on a logarithmic scale; the horizontal dashed line indicates the median depth of 882.5 analyses, and alternating grey shading groups subtypes by cancer root. In the lower track, aligned stacked bars show the proportions of defining analyses from each public data source (GENIE, GEO, cBioPortal, TCGA and Other (literature-curated public studies)).

The overall atlas construction workflow is summarized in Figure 2A. Within each root, analyses were represented in a normalized gene-level CNV comparison space. Candidate graph and clustering configurations spanning combinations of distance metric, retained principal components, *K*-nearest-neighbour (KNN) neighbourhood size and Leiden resolution were evaluated using graph topology, clustering stability and platform-association criteria to identify an initial community structure. Initial communities were subsequently refined through 20 subsampling repeats. Neighbouring communities that repeatedly clustered together were considered for merging, and boundaries with reproducible chromosome-scale or gene-level differences were retained. The refined communities were annotated using chromosome-scale gain and loss patterns (Track 1) together with subtype-associated gene-level CNV signals in the gain, high-level gain, loss and high-level loss channels (Track 2). Core communities were identified through discovery stability, whereas additional communities satisfying the conditional-selection procedure were retained as a secondary reference layer.

**Figure 2.**
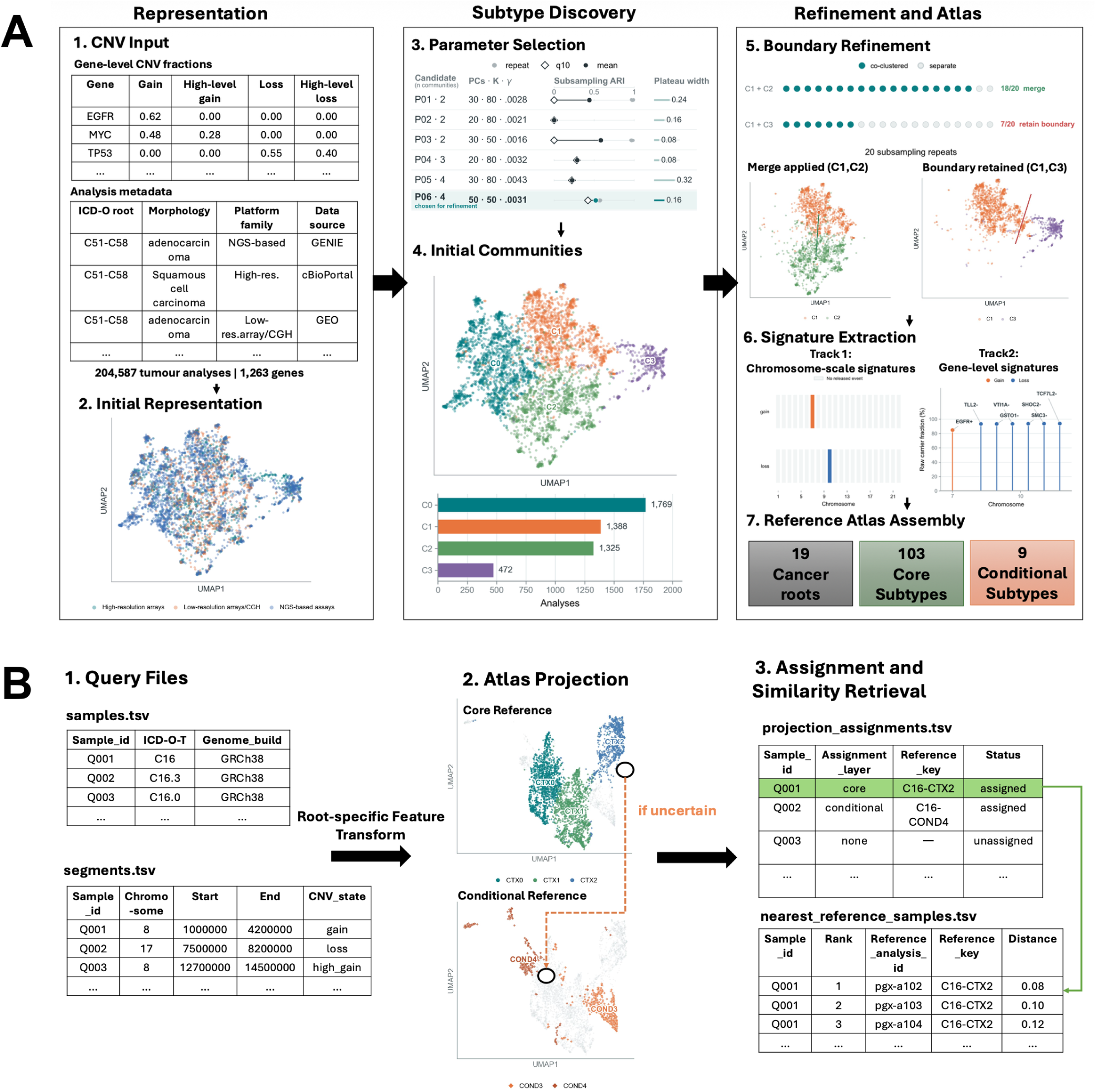
Construction and application of the CNVSIM reference atlas. **(A)** Root-wise construction of the CNVSIM reference atlas. Gene-level CNV profiles and harmonized sample metadata were represented within each atlas root before graph-based subtype discovery. Candidate graph configurations were evaluated using graph topology, clustering stability and platform-association criteria, with the liver/biliary root shown as an example. Initial communities were refined across 20 subsampling repeats by merging consistently co-clustering communities while retaining reproducible subtype boundaries. Separate core and conditional selection procedures retained 103 core and nine conditional reference subtypes across 19 atlas roots. Chromosome-scale signatures (Track 1) and subtype-associated gene-level signatures (Track 2) were used to annotate the retained subtypes. **(B)** Projection of new CNV profiles into the reference atlas. Query CNV profiles are represented in the corresponding root-specific comparison space and evaluated hierarchically against the core reference layer and, when necessary, eligible conditional subtypes. The stomach root illustrates the two reference layers in UMAP space. Projection returns either a calibrated subtype assignment or an explicit no-assignment status together with ranked nearest reference profiles.

Together, these formed a reference atlas comprising 112 CNV-defined subtypes across 19 cancer roots, including 103 core and nine conditional subtypes. The released atlas was defined by 153,033 analyses supporting the core reference layer and 2,848 supporting the conditional layer. Cross-study integration provided substantial reference depth. Subtypes were defined by 206 to 14,346 analyses (median, 882.5; Figure 1B). All subtypes integrated profiles across all three assay-platform families and multiple different data sources, including GENIE, GEO, cBioPortal, TCGA and other curated studies (Figure 1B). Each subtype was assigned a stable identifier together with diagnostic, assay-platform and data-source annotations, providing a fixed reference for downstream projection. New CNV profiles, including those from cell lines, can be projected into the reference atlas (Figure 2B). Queries are first compared with the core layer; eligible conditional subtypes are evaluated only when the core result is ambiguous or unresolved. Projection returns either a calibrated subtype assignment or an explicit no-assignment status, together with the nearest reference analyses. For assigned queries, CNVSIM further retrieves chromosome-scale and gene-level CNV signatures, available within-diagnosis somatic mutation associations, and candidate cell-line models with CNV signature evidence.

### Chromosome-scale and gene-level signatures characterize CNV reference subtypes

CNVSIM characterized the released subtypes using two complementary signature tracks to describe their CNV architecture relative to other analyses in the same anatomical root (Figure 3). Track 1 captured 295 subtype-enriched chromosome-scale gain and loss events. Track 2 identified 4,081 subtype-enriched gene-level CNV events. Together, these tracks annotated 108 of the 112 subtypes: 71 by both tracks, 18 by Track 1 alone and 19 by Track 2 alone (Table S1). The remaining four core subtypes were reproducibly recovered as communities but had no released signature events.

**Figure 3.**
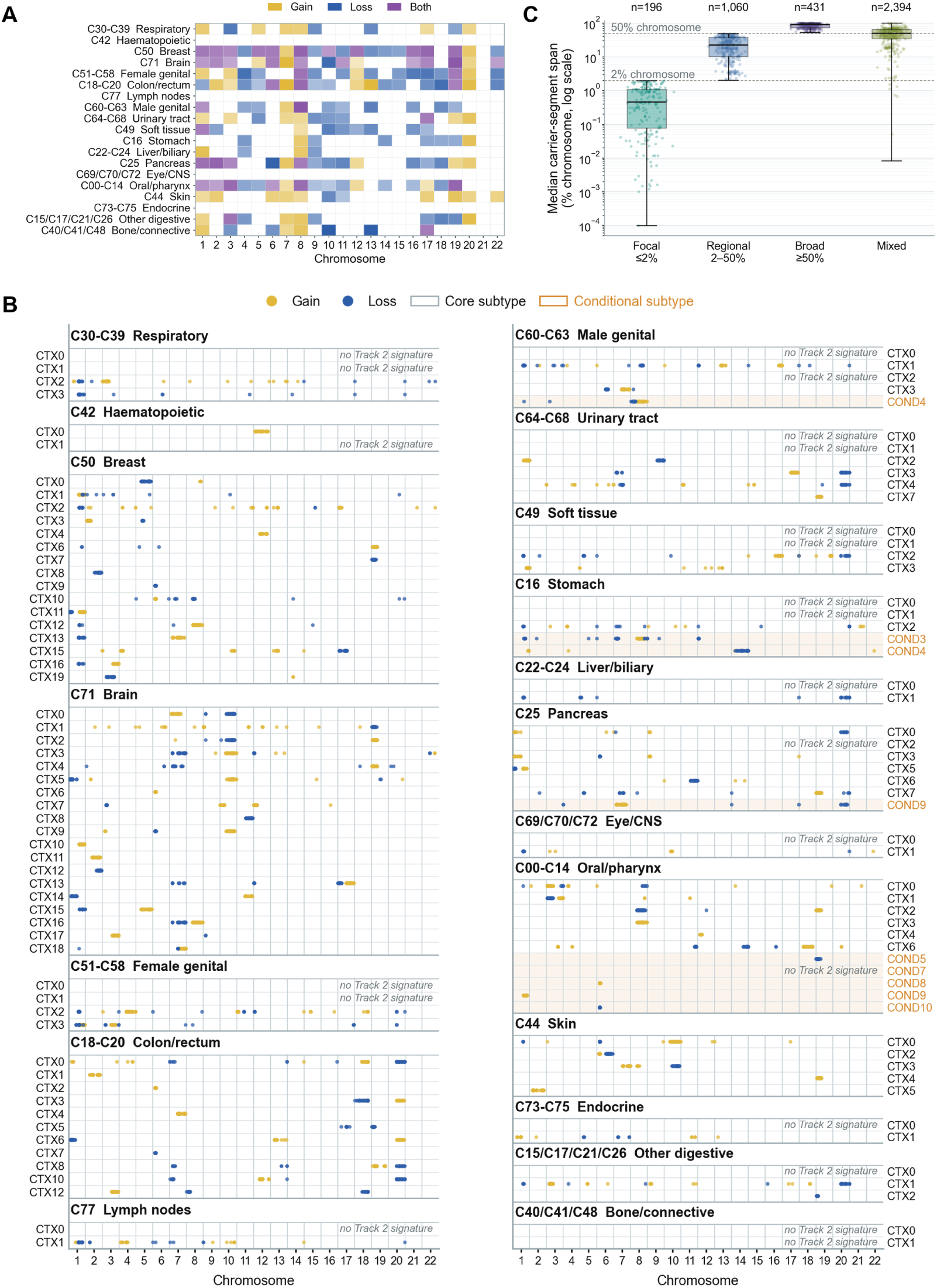
Chromosome-scale and gene-level CNV signature landscape. **(A)** Track 1 chromosome-scale gain and loss patterns across the 19 atlas roots. Colour encodes event direction (gain, loss, or both). Shading reflects carrier prevalence for selected events, weighted by subtype size relative to the total number of defining analyses in each root. **(B)** Genomic distribution of the 4,081 released Track 2 gene-level signature events across the 112 reference subtypes. Each row represents a core (CTX) or conditional (COND) subtype, and each point marks a selected gene-level CNV signature event at its genomic position on autosomes 1–22; subtypes without released Track 2 events are indicated. Gain and loss denote the combined all- and high-level gain or loss channels, with relative enrichment assessed using within-analysis mean-centred CNV fractions. **(C)** Segment-span distribution of the 4,081 released Track 2 signature events. The y-axis shows the median carrier-segment span as a percentage of chromosome length on a logarithmic scale, with horizontal dashed lines marking the 2% and 50% thresholds. Events are classified as focal (at most 2%), regional (above 2% and below 50%), broad (at least 50%), or mixed (at least two span categories each accounting for at least 25% of carrier analyses). Event counts are shown above each group.

Figure 3A summarizes the root-level distribution of Track 1 signatures. Track 1 captured several well-established chromosome-scale CNV patterns. Brain subtypes included the characteristic combination of chromosome 7 gain and chromosome 10 loss observed in glioblastoma^25^. Colon and rectum subtypes included gains involving chromosomes 13 and 20 together with chromosome 18 loss, consistent with recurrent colorectal cancer CNV^26^. Breast subtypes displayed extensive but subtype-specific combinations of chromosome-scale gains and losses, in line with the copy-number heterogeneity underlying established breast cancer genomic groups^17^. Several chromosome-scale signatures also recurred across anatomical roots and morphological diagnoses. Among the core subtypes, 23 chromosome-scale gain or loss signatures were selected in at least five cancer roots. Gains involving chromosome 8, predominantly affecting the long arm (8q), were the most widespread and occurred in 18 core subtypes across 14 roots, spanning subtypes dominated by adenocarcinoma, infiltrating duct carcinoma or squamous cell carcinoma. Gains involving chromosome 1 (predominantly 1q) occurred in subtypes from 13 roots, and gains involving chromosome 7 in subtypes from 11 roots. Subtypes with chromosome 7 gain included those dominated by glioblastoma or adenocarcinoma.

Figure 3B summarizes the genomic distribution of Track 2 events. Track 2 assessed relative enrichment using within-analysis mean-centred CNV fractions across the gain, high-level gain, loss and high-level loss channels, with unadjusted subtype and background frequencies reported alongside these statistics to aid interpretation. The released events involved 1,045 of the 1,263 cancer-associated genes and added locus-level detail to chromosome-scale patterns; for example, the glioblastoma-predominant subtype C71-CTX0 combined chromosome 7 gain and chromosome 10 loss with *EGFR* gain, *PTEN* loss and high-level *CDKN2A* loss.

A gene-level CNV signal alone does not indicate whether a locus is altered by a narrow focal event or carried within a larger segment spanning many genes. To provide this structural context, we next examined the spans of the original CNV segments supporting each Track 2 event (Figure 3C). Only 196 events (4.8%) were predominantly supported by focal segments spanning at most 2% of the chromosome. A further 1,060 events (26.0%) were classified as regional, with supporting segments spanning more than 2% but less than 50% of the chromosome, whereas 431 (10.6%) were classified as broad, with spans of at least 50%. The largest class comprised 2,394 mixed events (58.7%), for which at least two of the focal, regional and broad segment categories each accounted for at least 25% of carrier analyses. The resulting focal, regional, broad or mixed segment-span annotation is provided for every released Track 2 event.

To determine whether these mixed segment-span patterns arose from pooling data across different assay platforms, we repeated the classification separately for each platform family and individual assay platform (Table S2). Among the 2,394 mixed events, 1,775 (74.1%) had at least 10 segment carriers in each of two or more platform families and were included in the family-level assessment. Of these, 1,767 (99.5%) remained mixed within at least one family. At the finest available assay-platform level, 1,790 events (74.8%) met the same carrier requirement, and 1,761 (98.4%) remained mixed within at least one platform. These results show that pooling profiles across assay platforms was insufficient to explain the mixed segment-span patterns. The remaining variation may reflect a combination of biological differences in CNV boundaries and technical variation within individual platforms.

Together, the two signature tracks describe subtype-associated chromosome-scale gain and loss architecture and relative CNV differences at cancer-associated gene loci. Segment-span annotations indicate whether each gene-level event is predominantly focal or forms part of a larger or variably sized alteration. These annotations connect subtype labels to explicit CNV features, providing a structural basis for interpretation and comparison.

### CNV subtypes complement recorded diagnoses and show within-diagnosis somatic mutation associations

The number of released subtypes ranged from two to 19 per anatomical root (Figure 4A), and these CNV-defined subtypes did not map one-to-one onto recorded diagnostic labels. A common diagnosis could span several CNV architectures: infiltrating duct carcinoma was the dominant morphology in all 16 breast subtypes (73.7–89.0% of defining analyses), and colon adenocarcinoma was the leading NCI Thesaurus (NCIt) diagnosis in all 11 colon and rectum subtypes. Conversely, a single subtype could combine several diagnostic labels: the dominant morphology accounted for only 16.3–25.5% of defining analyses in the two haematopoietic subtypes and 26.4–34.2% in the two lymph-node subtypes, and the dominant NCIt diagnosis accounted for a median of 43.9% across all 112 subtypes (Table S1). By linking these diagnostic subdivisions and groupings to explicit chromosome- and gene-level CNV signatures, CNVSIM makes CNV heterogeneity within and across diagnostic labels directly interpretable.

**Figure 4.**
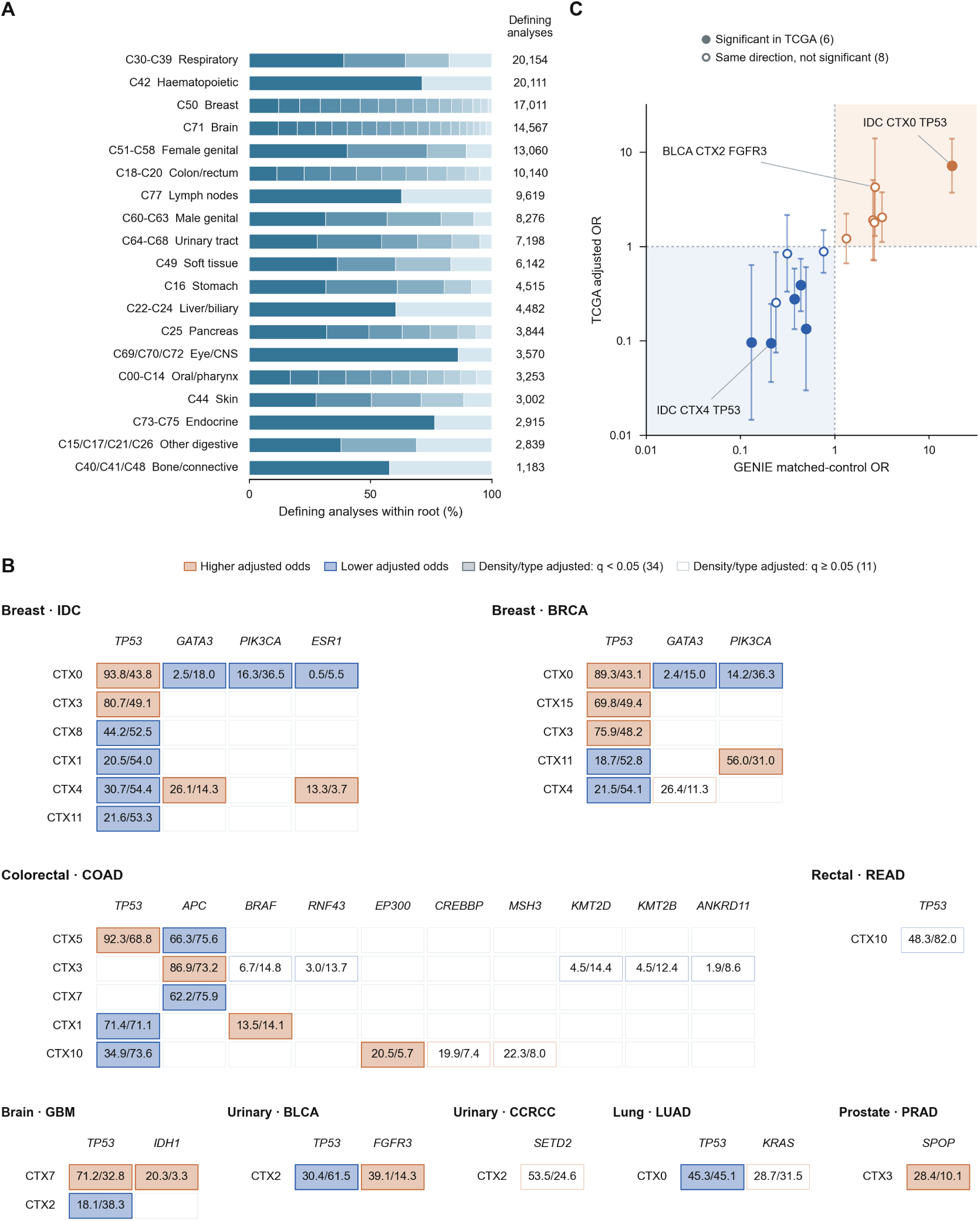
Reference subtype composition and within-diagnosis somatic mutation associations. **(A)** Distribution of the 155,881 defining analyses across 112 reference subtypes within 19 anatomical roots. Each bar sums to 100%, with segments representing subtypes ordered from largest to smallest within each root. The total number of defining analyses in each root is shown on the right. **(B)** The 45 within-diagnosis subtype–gene mutation associations identified in GENIE across three CNV-adjustment models (*q <* 0.05). Within each diagnosis block, rows represent subtypes and columns represent genes. Cell numbers show unadjusted mutation frequencies in the target subtype and its same-diagnosis controls (target/control, %), while orange and blue indicate higher and lower adjusted mutation odds in the target subtype. Filled cells (*n* = 34) remained significant after further adjustment for mutation density and primary versus metastatic status, outlined cells with numbers (*n* = 11) mark the remaining selected associations. **(C)** Cross-cohort comparison of the 14 associations that could also be evaluated in TCGA. Points compare the adjusted odds ratio (OR) in GENIE (x-axis) with the corresponding adjusted OR and 95% confidence interval in TCGA (y-axis) on logarithmic scales. Dashed lines mark OR = 1, and shaded quadrants indicate the same effect direction in both cohorts. Filled circles (*n* = 6) were significant across all three models in TCGA (*q <* 0.05), and open circles (*n* = 8) agreed in direction without reaching significance (Table S5).

Some CNV subtypes also differed in their somatic mutation patterns within a shared diagnosis, although mutation data were not used to define the subtypes. Across 40,254 evaluable GENIE analyses, 45 associations were identified in 17 of the 81 subtypes with comparisons estimable in all three models across six anatomical roots and 18 genes (Figure 4B). Each comparison tested a single gene, contrasting a target subtype with other eligible subtypes sharing the same anatomical root and recorded OncoTree diagnosis under three adjustment models. All three models adjusted for genome-wide CNV burden, local gene copy number, and assay panel. For each selected association, all three models agreed on whether mutation odds were higher or lower in the target subtype than in its controls. All 45 associations were significant in every model after correction for multiple testing. Thirty-four of the 45 associations remained significant after further adjustment for reported mutation density and whether the sample came from a primary or metastatic tumour (Table S5).

*TP53* accounted for 19 of the 45 associations across five anatomical roots, with mutation frequencies higher in some subtypes and lower in others within the same diagnosis. In invasive ductal carcinoma (IDC), C50-CTX0 showed a higher *TP53* mutation frequency (93.8% versus 43.8%) and lower *GATA3* (2.5% versus 18.0%) and *PIK3CA* (16.3% versus 36.5%) frequencies than its same-diagnosis controls. By contrast, C50-CTX4 showed the opposite pattern, with a lower *TP53* frequency (30.7% versus 54.4%) and higher *GATA3* frequency (26.1% versus 14.3%) relative to its own controls (Figure 4B; Table S5). Outside breast cancer, bladder urothelial carcinoma analyses in C64_C68-CTX2 showed higher *FGFR3* (39.1% versus 14.3%) and lower *TP53* (30.4% versus 61.5%) mutation frequencies, and prostate adenocarcinoma analyses in C60_C63-CTX3 showed a higher *SPOP* mutation frequency (28.4% versus 10.1%).

Some of these associations were also supported in TCGA, a separate cohort with exome-based rather than panel-based mutation data. Fourteen of the 45 associations could be compared in TCGA, and all 14 showed the same direction in both cohorts. Six were also statistically significant, all in IDC, which accounted for nine of the 14 comparisons; these included the contrasting *TP53* associations in C50-CTX0 and C50-CTX4 (Figure 4C; Table S5).

Overall, CNVSIM provides a structural organization complementary to pathological diagnosis, helping to interpret the CNV heterogeneity observed within and across diagnostic labels. Where paired mutation data and within-diagnosis sample sizes permitted, some of these CNV-defined subtypes also showed somatic mutation associations, with cross-cohort support for selected associations, linking part of this structural heterogeneity to accompanying mutational differences. The representative subtypes examined next illustrate how these diagnostic, structural, and mutational patterns come together in individual cancer roots.

### Representative subtypes illustrate CNV structure in diagnostic and somatic mutation contexts

Representative examples from the breast, colon and rectum, and brain roots illustrate how combined annotations connect reference subtypes to established cancer biology (Figure 5). For each subtype, diagnostic composition is shown together with its Track 1 chromosome-scale signature, Track 2 gene-level events and selected mutation associations.

**Figure 5.**
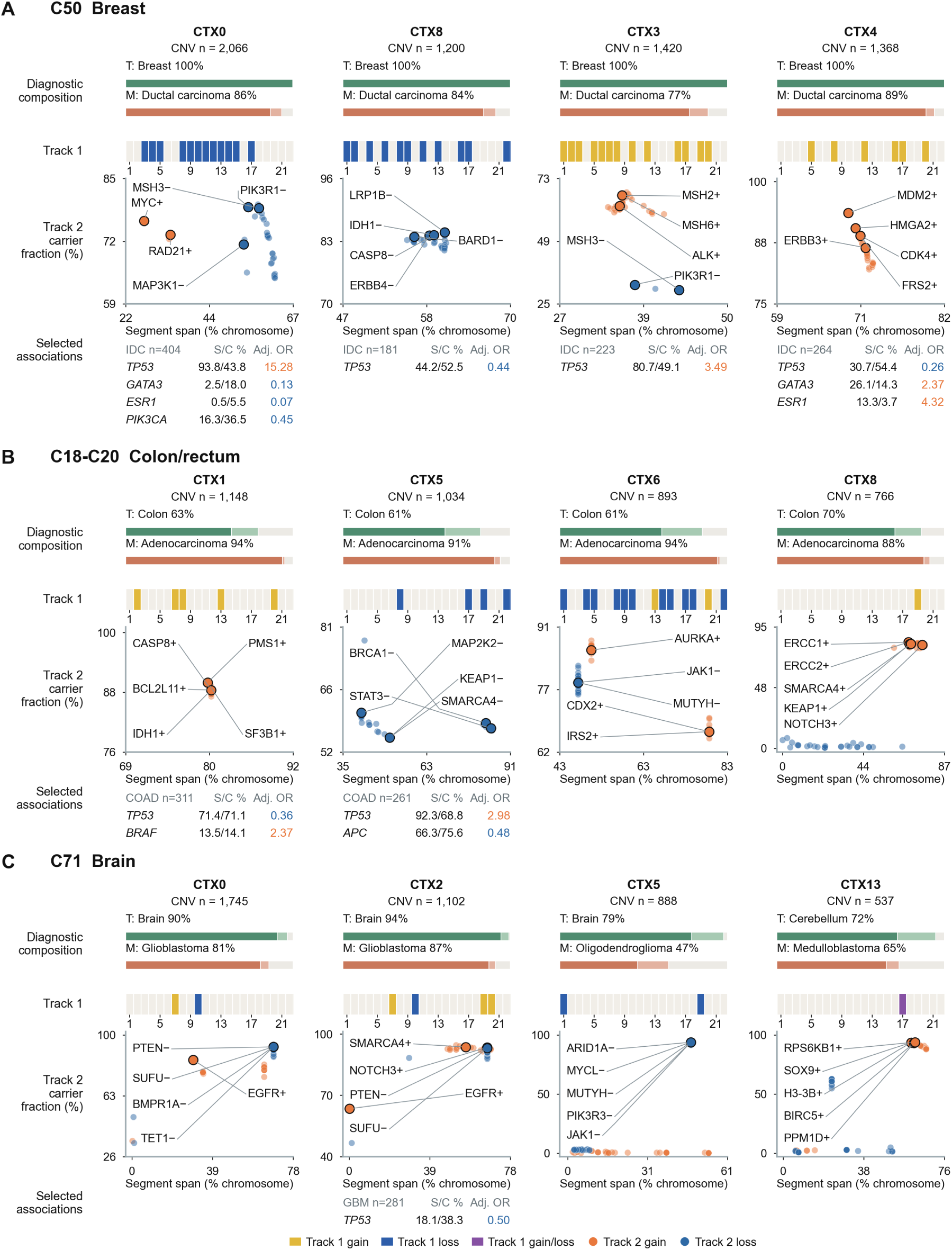
Representative subtypes across diagnostic and somatic mutation contexts. Representative released subtypes from the **(A)** C50 breast root, **(B)** C18–C20 colon/rectum root, and **(C)** C71 brain root. Each column represents one subtype and reports its number of defining CNV analyses (*n*). Diagnostic composition bars show the most frequent fine-grained ICD-O topography (T) and morphology (M) labels. Track 1 strips show enriched chromosome-scale gain, loss, or gain/loss events across autosomes 1–22. Track 2 plots show each gene-level event by its median carrier-segment span (% chromosome) on the x-axis and the percentage of subtype-defining analyses with supporting segments on the y-axis; representative genes are annotated with + or −, denoting a gain or loss event. Where present, tables at the bottom report selected within-diagnosis somatic mutation associations in GENIE for the indicated OncoTree diagnosis, showing unadjusted mutation frequencies in the subtype and its same-diagnosis controls (S/C %) and the primary adjusted odds ratio (Adjusted OR, coloured orange for higher and blue for lower adjusted odds).

The breast examples linked distinct chromosomal regions and mutation patterns within infiltrating duct carcinoma (77.4–89.0% of defining analyses) to recognized breast cancer genomic features (Figure 5A). Among subtypes with chromosome-scale loss signatures, CTX0 combined losses led by chromosomes 5, 4 and 14, including *PIK3R1* and *MAP3K1* on chromosome 5, with frequent *TP53* mutation and infrequent *GATA3* and *PIK3CA* mutation in invasive ductal carcinoma, a profile resembling features reported for basal-like breast cancer^27^. By contrast, CTX8 had losses led by chromosomes 2 and 13, including *BARD1* and *ERBB4* on chromosome 2, and a lower *TP53* mutation frequency than its same-diagnosis controls (44.2% versus 52.5%; Table S5). Among gain-dominated subtypes, CTX3 carried widespread gains, including *ALK*, *MSH2* and *MSH6* on chromosome 2, and showed a higher *TP53* mutation frequency (80.7% versus 49.1%), whereas CTX4 combined gains of chromosomes 12 and 20, including *MDM2* and *CDK4* on chromosome 12, with lower *TP53* and higher *GATA3* mutation frequencies. *MDM2* amplification is among the p53-pathway alterations reported in breast cancer, particularly in Luminal B tumours^27^.

Colon and rectum subtypes with similar diagnostic composition showed different combinations of chromosome-scale CNV signatures (Figure 5B). All four displayed subtypes were dominated by adenocarcinoma morphology (87.9–94.1%) and colon topography (60.9–70.4%). CTX1 was gain-dominated, with gains of chromosomes 2, 7, 8, 13 and 20. CTX6 shared the chromosome 13 and 20 gains but combined them with multiple chromosome-scale losses; its gained regions included *CDX2* and *IRS2* on chromosome 13 and *AURKA* on chromosome 20. Previous studies identified amplified *CDX2* as a colorectal lineage-survival oncogene and reported amplification of the mitotic kinase gene *AURKA* in colorectal cancer^28,29^. CTX8 instead had a restricted chromosome 19 gain signature, accompanied by gene-level gains involving *ERCC1*, *ERCC2*, *SMARCA4* and *KEAP1*. For these gene-level gains in CTX6 and CTX8, median supporting-segment spans covered at least half of the chromosome, placing the annotated loci within larger CNV regions. CTX5 showed a contrasting loss signature involving chromosomes 8, 17, 19 and 22. Within colon adenocarcinoma, it also showed higher *TP53* (92.3% versus 68.8%) and lower *APC* (66.3% versus 75.6%) mutation frequencies than its same-diagnosis controls (Table S5); *TP53* lies on chromosome 17, which is recurrently deleted in colorectal cancer^30^.

The brain examples showed both correspondence with recognized diagnostic entities and additional variation within them (Figure 5C). CTX0 and CTX2 were both dominated by glioblastoma, accounting for 80.6% and 87.1% of their defining analyses. Both shared the chromosome 7 gain and chromosome 10 loss pattern^25^, but CTX2 additionally showed gains of chromosomes 19 and 20, including gene-level gains involving *SMARCA4* and *NOTCH3* on chromosome 19. Concurrent gain of chromosomes 19 and 20 has been described in a subgroup of *IDH*-wild-type glioblastoma^31^. Within glioblastoma, CTX2 also showed a lower *TP53* mutation frequency than its same-diagnosis controls (18.1% versus 38.3%; Table S5). CTX5 had oligodendroglioma as its largest morphology group (46.5%) and showed chromosome 1 and 19 losses; its gene-level losses on chromosome 1, including *ARID1A* and *MYCL*, had median supporting-segment spans of about half the chromosome, a pattern compatible with the 1p/19q loss architecture central to the molecular classification of oligodendroglial tumours^32^. CTX13 was dominated by medulloblastoma and cerebellar topography and showed both gain and loss signatures involving chromosome 17, consistent with the recurrent chromosome 17 imbalance reported across medulloblastoma groups^33^.

Together, these examples show that the diagnostic subdivisions described above can be read in biological terms: subtypes sharing a diagnosis differed in specific chromosomes and genes, and several resembled known cancer features. Cross-study integration provided the reference depth supporting this subtype resolution: TCGA contributed fewer than 200 defining analyses to 101 of the 112 subtypes, whereas every subtype was defined by at least 206 analyses across sources. Because these annotations are attached to fixed subtype identifiers, a new cancer or cell-line profile assigned to a subtype can be interpreted against the same structural patterns, diagnostic composition, and subtype-level mutation associations across studies, rather than analyzed in isolation. We next tested how far this reference could be reproduced and how reliably it assigned new profiles.

### Reconstruction and projection tests characterize reference reproducibility

Every released subtype drew on at least ten assay platforms and four data-source groups, with no single platform or source contributing 75% or more of its defining analyses. To assess CN-VSIM for downstream applications, we examined whether core subtype definitions and signatures were retained after data reduction, whether held-out profiles from represented studies could be assigned consistently to fixed subtype labels, and whether profiles from an excluded cohort could be assigned to references rebuilt without that cohort (Figure 6).

**Figure 6.**
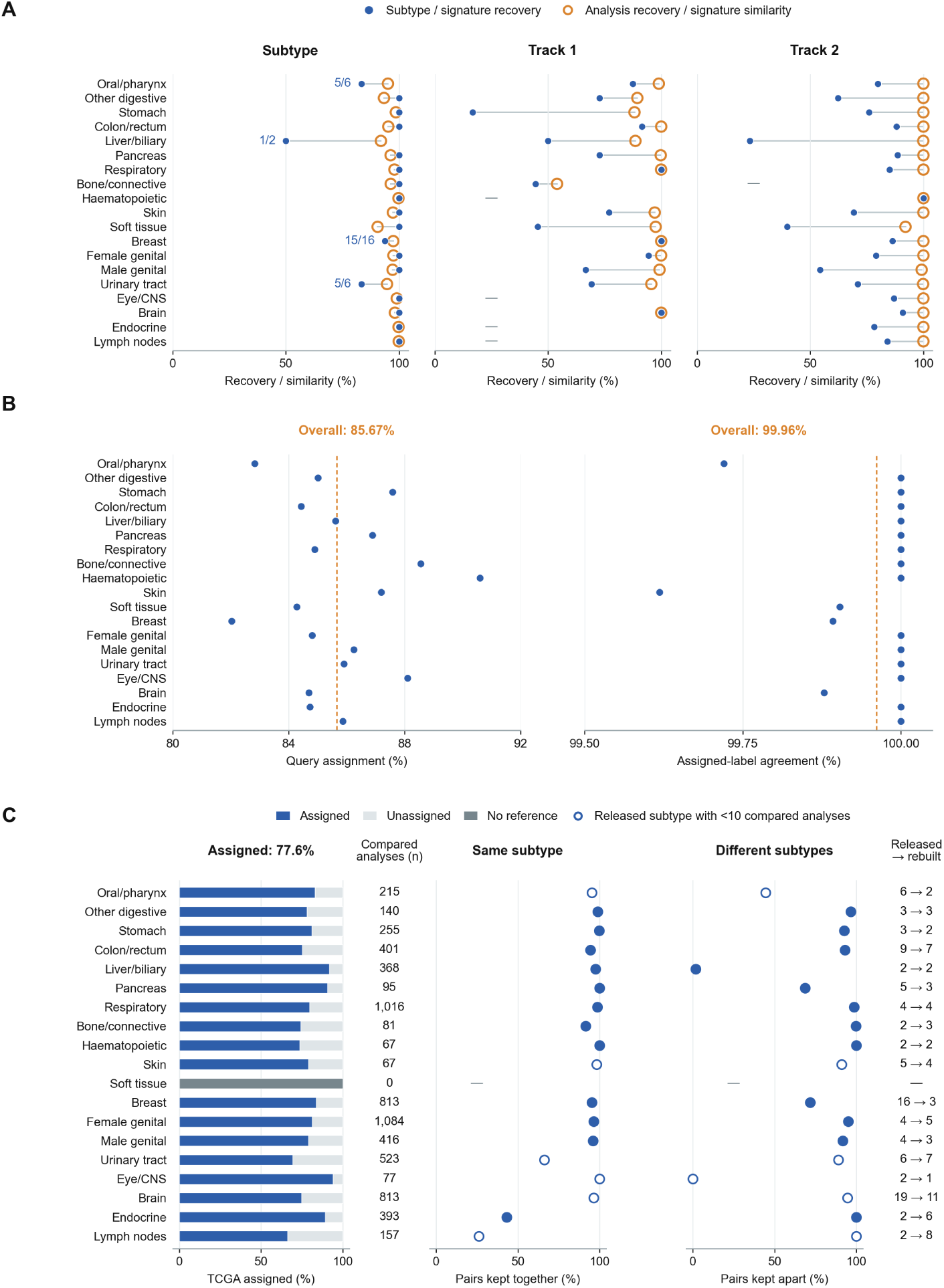
Reproducibility of reference subtypes, signatures and query projection. **(A)** Recovery of the 103 core subtypes and their signatures in five rebuilds, each withholding 20% of analyses within each root’s fixed comparison space while retaining the released profile coordinates and parameters. Left, filled blue circles show the minimum fraction of core subtypes recovered across rebuilds; open orange circles show the median fraction of retained defining analyses in communities linked to their original subtype. Annotated fractions give minimum-fold recovered/total subtype counts. For Track 1 and Track 2, blue circles show median recovery of released signature events with the same CNV state and enrichment direction, and orange circles show median enrichment-profile cosine similarity. **(B)** A projection test using 30,609 held-out core defining analyses and 122,424 core analyses in the projection reference. The left panel shows assignment coverage and the right panel assignment accuracy among assigned queries, defined as agreement with their original atlas subtype labels. Orange dashed lines indicate the overall values (85.67% coverage and 99.96% accuracy). **(C)** Reference reconstruction without TCGA and projection of 10,156 held-out TCGA analyses. Left, percentages of held-out TCGA analyses assigned to a rebuilt subtype (blue; 77.6% overall), left unassigned (light grey), or in a root without a rebuilt reference (dark grey), followed by the number of analyses assigned under both references (*Compared analyses (n)*, total *n* = 6,981). Middle and right, percentages of same-subtype analysis pairs kept together and different-subtype pairs kept apart in the rebuilt reference, with the numbers of represented released and rebuilt groups (*Released* → *rebuilt*) shown on the right. Open circles indicate roots with fewer than 10 compared analyses in at least one represented released subtype.

To assess the reproducibility of the CNV signatures used for subtype interpretation, we repeated community discovery and signature extraction in five rebuilds, each withholding 20% of analyses within each root’s fixed comparison space while retaining the original profile coordinates and selected parameters (Figure 6A). The rebuilds recovered a median of 102 of the 103 core subtypes (minimum, 101). A median of 96.9% of retained core defining analyses belonged to rebuilt communities linked to their original subtype. Median recovery of individual signature events was 83.7% for Track 1 and 81.8% for Track 2, and global median cosine similarity of the enrichment profiles exceeded 0.999 for both tracks.

We next assessed whether query projection could link additional profiles consistently to the annotated reference. We withheld 30,609 core defining analyses as queries and constructed the projection reference from the remaining analyses, keeping the released subtype labels and previously selected root-specific parameters fixed (Figure 6B). Of the held-out queries, 26,223 (85.7%) met the assignment criteria, while the remainder were left unassigned. Among assigned queries, 26,213 (99.96%) matched their released subtype label, supporting consistent linkage to the existing reference annotations in this holdout setting.

To approximate how CNVSIM receives a cohort it has not seen, we removed all TCGA analyses before preprocessing, parameter selection and community discovery, rebuilt the references, and projected 10,156 held-out TCGA analyses (Figure 6C). The rebuilt references assigned 7,880 analyses (77.6%); 2,083 remained unassigned and 193 had no rebuilt subtype reference. Grouping correspondence was assessed for 6,981 TCGA analyses with subtype labels under both references. In 10 of the 18 comparable roots, at least 90% of same-subtype analysis pairs were kept together and at least 90% of different-subtype pairs were kept apart. Reconstruction also changed subtype resolution in some roots. The 16 represented breast subtypes mapped to three rebuilt groups: CTX0 and CTX8, which had chromosome-scale loss signatures, mapped mainly to one group, whereas CTX3 and CTX4, which had gain signatures, mapped mainly to another. The two lymph-node subtypes split into eight groups, while liver/biliary showed poor correspondence, with nearly all compared analyses assigned to one rebuilt group.

Together, these tests support the use of CNVSIM as a fixed reference for assigning and interpreting CNV profiles. In the internal holdout test, 85.7% of queries were assigned and 99.96% of those assignments matched released labels; the TCGA-excluded references assigned 77.6% of the held-out cohort. The fixed-setting rebuilds showed that the released core structure was largely retained after data reduction. After TCGA exclusion, at least 90% of same-subtype analysis pairs remained together in 15 of the 18 comparable roots, although subtype boundaries varied under the changed reference data and reselected parameters. The versioned subtype definitions provide a common basis for comparing assigned profiles across studies.

### Reference projection identifies candidate cell-line models and gaps in CNV representation

Cancer cell lines are widely used as experimental models in biomedical research and pharmacological screening. Matching their CNV profiles to annotated subtypes in the atlas helps prioritize models for experiments focused on specific tumour CNV patterns. We projected 4,605 CNV profiles from 1,704 Cellosaurus-linked cell lines^34^ into the atlas within their anatomical roots, identifying 1,099 recommended cell-line–subtype matches. Of these, 413 were supported by multiple profiles and 686 by a single eligible profile.

Recommended models were available for 90 of the 112 released reference subtypes (Figure 7A; Table S4). Sixteen subtypes had no calibrated cell-line match, and six had matched models that did not meet the recommendation criteria. For example, the prostate subtype C60_C63-COND4 was defined by 604 analyses, 94.2% of which had an adenocarcinoma morphology yet had no calibrated cell-line match. Its chromosome 8 gain and loss signatures, including *MYC* gain and *NKX3-1* loss, were compatible with the 8p-loss/8q-gain pattern reported in prostate cancer^35^. Likewise, the breast subtype C50-CTX12 had no calibrated cell-line match despite being defined by 764 analyses and carrying a chromosome 8 gain pattern involving *MYC*, *RAD21* and other chromosome 8 genes, consistent with previously described breast cancer genomic patterns^36^. These examples show that a subtype can be well represented in the atlas and carry a recognized CNV pattern yet lack a recommended cell-line model.

**Figure 7.**
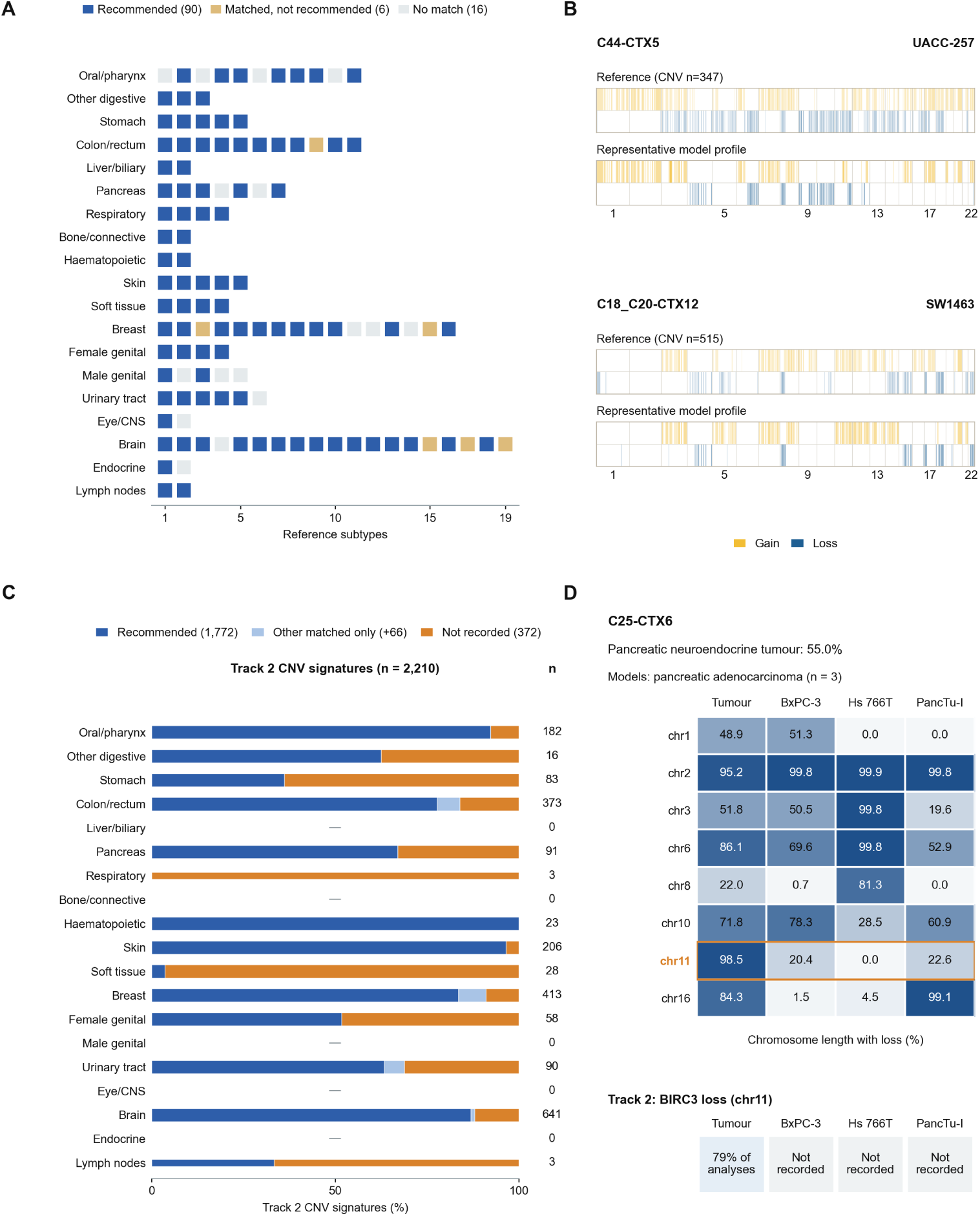
Candidate cell-line models and signature coverage across reference subtypes. **(A)** Availability of Cellosaurus-linked cell-line models across the 112 released reference subtypes. Each square represents one subtype grouped by anatomical root. Blue indicates subtypes with at least one recommended model (*n* = 90), gold indicates subtypes with matched models that did not meet the recommendation criteria (*n* = 6), and light grey indicates subtypes with no calibrated cell-line match (*n* = 16). **(B)** Representative subtype–cell-line matches for the melanoma subtype C44-CTX5 (UACC-257; CVCL_1779) and the colorectal subtype C18_C20-CTX12 (SW1463; CVCL_1718). Reference subtype profiles show mean encoded gain and loss scores (scaled from 0 to 1) across subtype-defining analyses, and cell-line profiles show the representative recommended profile. **(C)** Cell-line coverage of 2,210 Track 2 signatures with higher unadjusted subtype frequencies than root background, restricted to subtypes with recommended models. Dark blue, recorded in at least one recommended model (*n* = 1,772); light blue, recorded only in other matched models (*n* = 66); orange, not recorded in any matched model of that subtype (*n* = 372). Signature counts are shown at right. **(D)** Diagnostic context and CNV comparison for pancreatic subtype C25-CTX6 and its recommended models. Pan-creatic neuroendocrine tumour accounts for 55.0% of defining analyses, whereas BxPC-3, Hs 766T and PancTu-I are annotated as pancreatic adenocarcinoma. Top, loss coverage for the eight Track 1 signature chromosomes, shown as the median percentage of chromosome length across defining tumour analyses and the value for each model profile; chromosome 11 is highlighted in orange. Bottom, *BIRC3* loss, recorded in 79% of defining analyses and in none of the three models.

To illustrate what a recommended match represents, we compared two reference subtype profiles with their recommended cell lines (Figure 7B). UACC-257 was a match supported by multiple profiles for the melanoma subtype C44-CTX5, and SW1463 for the colorectal subtype C18_C20-CTX12. In both examples, the recommended cell lines shared major components of the gain and loss architecture observed across the corresponding reference subtype while retaining expected cell-line-specific differences.

We next examined how the matched models represented the gene-level CNV signatures of their assigned subtypes. Among 2,210 Track 2 signature annotations with higher unadjusted subtype frequencies than root background, restricted to subtypes with recommended models, 1,772 (80.2%) were recorded in at least one recommended model, 66 (3.0%) only in other matched models, and 372 (16.8%) in no matched model of that subtype (Figure 7C). For example, in the pancreatic subtype C25-CTX6, pancreatic neuroendocrine tumour accounted for 55.0% of defining analyses, whereas its three recommended models, BxPC-3, Hs 766T and PancTu-I, were annotated as pancreatic adenocarcinoma (Figure 7D). Chromosome 11 loss covered a median of 98.5% of the chromosome across defining analyses compared with 0–22.6% in the three model profiles, and *BIRC3* loss on chromosome 11 was recorded in 79.0% of defining analyses but in none of these models. The catalogue also included QGP-1, annotated as pancreatic somatostatinoma, which was assigned to C25-CTX2 and whose single available profile recorded neither chromosome 11 loss nor *BIRC3* loss. Thus, none of the four evaluated profiles reproduced the extensive chromosome 11 loss pattern of C25-CTX6.

The released CNVSIM atlas provides a reusable reference for CNV profile comparison and experimental-model selection. Users can project new CNV profiles, including those from cell lines, to obtain calibrated subtype assignments together with the nearest reference profiles. They can also retrieve recommended cell-line models for a reference subtype or look up the CNV-defined subtype assigned to a particular Cellosaurus cell line. These projection and model-selection tools are available through the CNVSIM GitHub repository (Figure S1).

## DISCUSSION

CNVSIM establishes an interpretable cross-study reference atlas for genome-wide CNV profiles. Previous studies demonstrated that CNV profiles capture biologically meaningful structure in cancer genomes, whether by identifying recurrent CNV landscapes, linking genome-wide CNV patterns to chromosomal-instability processes, identifying molecular groups within individual cancers and across pan-cancer multi-omic datasets, or by classifying samples into predefined diagnostic categories^3,13–18^. These approaches advanced understanding of CNV biology in cancer, and CNVSIM builds on them by organizing cross-study CNV structure into a fixed, versioned reference atlas that can be applied consistently to newly generated CNV profiles from individual samples, cohorts and experimental models.

Building such a reference from heterogeneous public CNV data requires priorities different from subtype discovery. Progenetix provides harmonized genome-wide CNV profiles together with diagnostic, assay, and study-provenance metadata, making cross-study reference construction at this scale possible^7^. Aggregation across public studies nevertheless retains both technical and biological heterogeneity. The objective of CNVSIM was to preserve recurrent biological variation while reducing the likelihood that technical variation determined subtype boundaries. Platform association was therefore considered when selecting candidate graph configurations, whereas diagnostic, assay platform and data source composition were retained as explicit annotations for every subtype, with every released subtype drawing on multiple platforms and source groups. These design choices cannot eliminate all residual platform or source effects, but they make the supporting evidence transparent and allow users to evaluate the biological interpretation and technical context of each subtype.

The value of such a reference ultimately depends on whether CNV captures biologically meaningful patterns of genomic variation. CNV is well suited to serve as a reference layer because it captures recurrent genome organization at a level between broad diagnostic classifications and individual gene-level alterations. Existing diagnostic classifications describe anatomical origin and histopathology but often encompass multiple recurrent CNV architectures, whereas individual gene-level events do not preserve the broader chromosomal context in which they occur. Consistent with previous molecular classifications of breast and colorectal cancer^17,37^, the atlas repeatedly resolved broad diagnostic entities into multiple CNV-defined reference subtypes. Conversely, individual CNV-defined subtypes frequently contained multiple morphology or disease labels within the same anatomical comparison space, indicating that recurrent CNV organization does not always follow existing diagnostic boundaries. Similar relationships have also been observed across cancer types, where recurrent chromosome-arm alterations extend beyond individual diagnostic categories^18,38^. Anatomical grouping therefore defines the comparison space, whereas CNV subtypes resolve recurrent genomic diversity within that space.

The two signature tracks make this organization interpretable by describing the CNV architecture associated with each reference subtype at chromosome and gene scales. Chromosome-scale signatures summarize recurrent total copy-number gain and loss architecture, whereas gene-level events localize cancer-associated loci within these broader genomic states^3,13^. Broad chromosomal alterations can reflect coordinated dosage effects across multiple genes and recurrent karyotypic states shaped during cancer evolution^39,40^. Most Track 2 events in the CN-VSIM atlas were supported by regional, broad or mixed segment spans rather than isolated focal alterations, indicating that subtype-associated gene-level events frequently form part of larger chromosomal changes. Although CNVSIM does not resolve allele-specific copy number or infer the mechanisms generating these alterations, it preserves the structural context needed to interpret subtype-associated gene-level CNV signals. Somatic mutations were not used to define subtypes, but their frequencies differed between some subtypes sharing a recorded diagnosis. In GENIE, these differences remained after adjustment for genome-wide CNV burden and local copy number. In TCGA, all 14 estimable comparisons agreed in direction with the GENIE estimates. These associations provide molecular context for part of the CNV heterogeneity captured by the atlas. Sequence-based and multi-omic classifications can further distinguish samples sharing similar CNV architecture, and CNVSIM is complementary to them^41–43^.

The reconstruction and projection tests support CNVSIM as a shared reference for interpreting additional CNV profiles. A query profile is projected within its anatomical root and, when assigned, linked to the annotations of a reference subtype and to candidate cell-line models with similar CNV architecture (Figure S1). Signature comparisons then show which alterations needed for an experiment are recorded in those candidates and where the evaluated catalogue leaves gaps. Earlier studies compared tumours and cell lines using genomic aberrations or multiomic profiles^44,45^. CNVSIM places this comparison within the same annotated reference used to interpret tumour profiles.

### Limitations of the study

The current atlas reflects the limitations of its source data and reference design. Public data remain unevenly distributed across cancer types, diagnostic groups, contributing studies, assay technologies and clinical settings. Some cancer roots therefore have substantially greater representation than others, which may affect the level of recurrent CNV structure that can be resolved. Chromosome-scale and segment-span annotations also depend on the resolution and genomic coverage of the original assays and on their segmentation procedures.

Subtype granularity depends on data composition and root-specific graph parameters. The released subtypes provide a fixed, versioned reference based on the best available settings among those evaluated for the current dataset under the selection criteria (STAR Methods; Table S3), rather than a unique clustering solution. Mutation associations were restricted to subtypes and diagnoses with sufficient matched data. Query projection requires the anatomical root of the input profile to be specified in advance. Cell-line recommendations are limited to models with eligible public CNV profiles. The absence of a recommendation identifies a gap in the evaluated catalogue rather than demonstrating that no suitable experimental model exists, while a recommendation indicates CNV similarity rather than complete genomic or functional equivalence.

### Future directions

Future development could extend both the coverage of the atlas and the evidence associated with its subtypes. Additional bulk CNV profiles could improve the representation of under-represented cancer roots and diagnostic groups, while compatible longitudinal and multi-region bulk profiles could be projected to compare relationships to reference subtypes across tumour regions, disease stages or treatment time points, although projection alone would not reconstruct evolutionary lineages or determine the temporal order of alterations^40^. Clone-level or pseudobulk profiles obtained from single-cell DNA sequencing or inferred from single-cell transcriptomes would require modality-specific preprocessing and calibration^46,47^. Additional molecular measurements, treatment responses and clinical outcomes could likewise be linked to projected CNV profiles without redefining the reference itself, and experimental systems such as organoids and patient-derived xenografts could be evaluated when suitable genome-wide CNV profiles become available and projection limits are validated for those models^48,49^.

### Conclusions

CNVSIM organizes heterogeneous public CNV profiles into an annotated reference atlas comprising 112 subtypes across 19 anatomical cancer roots. Its subtypes capture CNV diversity within recorded diagnoses and shared architectures across diagnostic labels, and some differ in somatic mutation frequency within a shared diagnosis. Calibrated projection supports interpretation of new profiles and selection of candidate cell lines based on CNV similarity. With fixed, versioned subtype definitions, CNVSIM provides not only a shared reference for CNV profile comparison, but also a foundation on which biological, clinical and experimental knowledge associated with recurrent CNV architecture can continue to grow.

## RESOURCE AVAILABILITY

### Lead contact

Requests for further information and resources should be directed to and will be fulfilled by the lead contact, Michael Baudis.

### Materials availability

This study did not generate new, unique reagents.

### Data and code availability

- Harmonized CNV profiles from neoplastic samples and cancer cell lines, together with associated metadata, were obtained from Progenetix (https://progenetix.org)^7^. The cancer-gene feature set was derived from OncoKB^9^ and the COSMIC Cancer Gene Census^10^, and cell-line identifiers were standardized using Cellosaurus^34^. Somatic mutation analyses used AACR Project GENIE release 17.0-public (https://genie.synapse.org/)^50^ and the public TCGA MC3 v0.2.8 callset (https://gdc.cancer.gov/about-data/ publications/mc3-2017)^51^; sample-level mutation records are available from the respective source repositories. The CNVSIM Atlas v1.0.1 data release, including reference subtype definitions and annotations, chromosome- and gene-level CNV signatures, the calibrated projection bundle, cell-line matching tables, and summary statistics from the validation and somatic mutation analyses, is available at Zenodo (https://doi.org/10.5281/zenodo.23982370) under the CC BY 4.0 license.
- CNVSIM source code is publicly available on GitHub (https://github.com/baudisgroup/ CNVSIM; release CNVSIM_v1.0.1) and archived at Zenodo (https://doi.org/10.5281/zenodo.23982370).
- Any additional information required to reanalyze the data reported in this paper is available from the lead contact upon request.

## ACKNOWLEDGMENTS

We thank Hangjia Zhao for the continued development, harmonization, and quality improvement of the Progenetix resource, whose sustained development made this work possible. We are grateful to Qingyao Huang, Paula Carrió-Cordó, Bo Gao, Rahel Paloots and other former members of the Baudis group for their contributions to the long-term development and curation of Progenetix. We especially thank Rahel Paloots for her work on the integration and annotation of cancer cell-line data, which enabled the experimental-model analyses presented in this study. Finally, we acknowledge the many investigators who generated and openly shared copy-number data through public repositories, making this reference atlas possible.

We acknowledge the American Association for Cancer Research for its financial and material support in developing the AACR Project GENIE registry. We thank the consortium members for their commitment to data sharing. Interpretations are the responsibility of study authors. The results reported here are also based in part on data generated by the TCGA Research Network (https://www.cancer.gov/tcga).

The project was supported by funding from the Swiss National Science Foundation (SNSF proposal 10.011.287).

## AUTHOR CONTRIBUTIONS

Conceptualization, J.Y. and M.B.; Methodology, J.Y.; Software, J.Y. and S.J.; Validation, J.Y.; Formal Analysis, J.Y. and J.B.-T.; Data Curation, J.Y.; Visualization, J.Y.; Writing – Original Draft, J.Y.; Writing – Review & Editing, J.Y. and M.B.; Resources, M.B.; Supervision, M.B.; Project Administration, J.Y. and M.B.; Funding Acquisition, M.B. All authors read and approved the final manuscript.

## DECLARATION OF INTERESTS

The authors declare no competing interests.

## DECLARATION OF GENERATIVE AI AND AI-ASSISTED TECHNOLOGIES

During the preparation of this work, J.Y. applied ChatGPT in order to edit language and improve clarity and organization. After using this service, all authors reviewed and edited the content as needed and take full responsibility for the content of the published article.

## SUPPLEMENTAL INFORMATION INDEX

**Table S1.** Diagnostic composition and CNV features of the released CNVSIM reference subtypes (Table_S1_subtype_root_features.xlsx). Related to Figures 3, 4 and 5. Reports the diagnostic, morphological, topographical, assay-platform and data-source composition of the 112 released reference subtypes, together with representative topography- and morphology-associated CNV features and recurrent chromosome-scale signatures across cancer roots.

**Table S2.** Platform-stratified assessment of mixed Track 2 segment-span events (Table_S2_ mixed_track2_platform_sensitivity.xlsx). Related to Figure 3. Reports platform-family and individual-platform assessments of the 2,394 mixed Track 2 events, including evaluability, persistence of mixed span classifications within platform groups, and Kruskal–Wallis comparisons of carrier-segment spans.

**Table S3.** Fixed analysis parameters, statistical models, and decision thresholds used in CN-VSIM (Table_S3_analysis_parameters.xlsx). Related to STAR Methods. Lists 158 parameters and decision rules across 19 analysis stages, including atlas construction, signature extraction, projection, mutation context, source holdout and cell-line evidence.

**Table S4.** Cell-line model availability, coverage gaps and Track 2 gene-event representation across CNVSIM subtypes (Table_S4_cell_line_coverage_and_gaps.xlsx). Related to Figure 7. Reports cell-line recommendation coverage across the 112 released subtypes and 438 Track 2 signature annotations with higher unadjusted subtype frequencies than root background that were not recorded in recommended cell-line models of the same subtype.

**Table S5.** Within-diagnosis somatic mutation associations in GENIE and cross-cohort comparisons in TCGA MC3 (Table_S5_mutation_context.xlsx). Related to Figure 4 and 5. Reports 45 GENIE associations after adjustment for genome-wide CNV burden and local copy number (34 retained after additional mutation-density and sample-type adjustment), 14 estimable TCGA MC3 comparisons, and representative subtypes with their CNV signatures and retained mutation associations.

**Figure S1.** CNVSIM application workflow (Figure_S1_use_case.pdf). Related to Figure 7. Worked example of CNV-profile retrieval through the public Progenetix API, projection-input preparation and integrated reference-context queries, together with Cellosaurus cell-line queries.

## STAR METHODS

### Method details

#### Data source and metadata harmonization

We used publicly available genome-wide CNV profiles (GRCh38) and associated metadata from Progenetix to construct and evaluate a reference atlas for cancer subtyping^7^. The source collection contained 257,365 analyses. Here, an analysis refers to a Progenetix analysis record; multiple analyses may originate from the same biosample. Sample eligibility for atlas evaluation was based on biosample-status annotations: “neoplastic sample”, “primary tumor sample”, “metastasis sample” or “recurrent tumor sample”. Cancer cell-line analyses were retained separately for downstream projection. Analyses annotated with the non-specific Experimental Factor Ontology (EFO) platform terms “DNA assay” (EFO:0001456) or “DNA array” (EFO:0002701) were excluded. The resulting dataset comprised 204,587 eligible neoplastic analyses and 4,605 eligible cancer cell-line analyses for downstream projection. Of the neoplastic analyses, 155,881 ultimately defined the 112 released reference subtypes: 153,033 in the core layer and 2,848 in the conditional layer.

We defined comparison roots using ICD-O-3 topography codes since they provide standardized, mutually exclusive anatomical site annotations across Progenetix that correspond largely to clinical ICD-10 malignancy categories. These topography annotations were mapped to 22 grouped anatomical roots, with closely related topography codes combined where necessary to maintain a sufficient number of analyses and anatomical coherence. These grouped roots defined the root-specific comparison spaces used throughout preprocessing, graph construction, and projection. C76 (ill-defined anatomical sites) and C80 (unknown primary sites) were excluded from subtype discovery. Among the remaining roots, C47 (Peripheral nerves and autonomic nervous system) did not yield a reproducible within-root subtype partition. The released atlas therefore comprised 19 root-specific comparison spaces. ICD-O topography, ICD-O morphology, and NCIt diagnosis annotations were retained for subtype description and downstream interpretation.

Assay-platform annotations were harmonized through a curated platform catalogue. Individual platform identifiers were assigned to three platform families: NGS-based assays, high-resolution arrays, and low-resolution arrays or CGH-based profiles. Array resolution was determined using existing platform annotations together with manual catalogue review, with a threshold of 180,000 probes used to distinguish high- from low-resolution arrays. Across the 204,587 analyses, 235 platform identifiers were represented, comprising 115,316 NGS-based analyses, 49,659 high-resolution arrays, and 39,612 low-resolution array or CGH-based profiles. Study provenance was summarized into GENIE, GEO, TCGA, cBioPortal, ArrayExpress, and other sources. Platform-family annotations were used during graph-parameter evaluation, whereas assay-platform and data-source composition were retained as annotations for each reference subtype.

#### CNV feature construction and data preprocessing

Gene-level CNV summaries were obtained from Progenetix for 1,325 cancer-associated genes defined as the union of OncoKB and the COSMIC Cancer Gene Census^9,10^. The frozen autosomal gene set and GRCh38 intervals are provided in projection_bundle/gene_intervals.tsv in the v1.0.1 data release. After restricting to autosomal genes, 1,263 genes were retained. For each analysis and gene, Progenetix records the fraction of the gene interval affected by gain, high-level gain, loss, and high-level loss.

Feature construction was performed independently within each grouped ICD-O-T root. For each analysis *s* and gene *g*, the four gene-level CNV fractions were converted into binary event indicators using a threshold of 0.10 for gain and loss channels and 0.20 for high-level gain and loss channels:

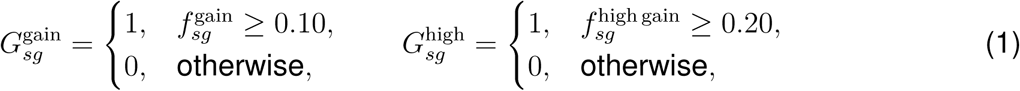

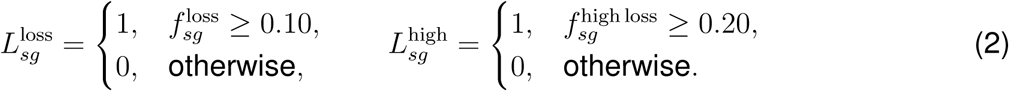

The paired indicators were then combined into direction-specific weighted discrete features:

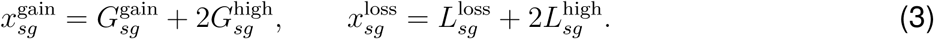

This encoding preserved CNV direction while assigning greater weight to high-level events. The gain and loss indicators were not mutually exclusive of their corresponding high-level indicators; a feature value of 3 therefore indicated that both thresholds were met. Each retained autosomal gene contributed one gain feature and one loss feature. Gene coverage was assessed separately within each root after excluding platform families represented by fewer than 50 analyses in that root. This prevented sparsely represented platform groups from determining gene-coverage estimates. Genes were required to have corresponding gene-level CNV records in at least 95% of analyses within every remaining platform family. All 1,263 autosomal genes satisfied this criterion, yielding 2,526 encoded features. Analyses without an encoded CNV event in any retained autosomal gene were excluded, leaving 174,670 analyses.

A preliminary quality-control procedure was then applied to identify small groups dominated by a single platform family. Within each root, encoded features were standardized to zero mean and unit variance, reduced to 50 principal components, projected with uniform manifold approximation and projection (UMAP; 30 neighbours, minimum distance 0.1, cosine distance, random seed 0), and clustered using hierarchical density-based spatial clustering of applications with noise (HDBSCAN; minimum cluster size 200; minimum samples 50). A non-noise cluster was removed only when it satisfied all three criteria: (i) it contained no more than 10% of the analyses in the root, (ii) at least 90% of its analyses originated from the same platform family, and (iii) its dominant ICD-O morphology accounted for less than 50% of the cluster. Across the 21 preprocessed anatomical roots (all grouped roots except C80, including C76 and C47 prior to final root selection), this procedure removed 3,135 analyses, retaining 171,535 analyses.

The retained analyses were then reprocessed to generate the final comparison space used for graph construction. Feature vectors were normalized to unit *L*_2_ norm, standardized by feature within each root, and represented by the first 50 principal components. *L*_2_ normalization reduced the influence of the total number of altered genes while preserving the relative gain and loss pattern of each profile. These root-specific principal-component coordinates formed the input for graph construction.

#### Reference atlas construction and subtype selection

Reference subtypes were identified independently within each root-specific comparison space using weighted *K*-nearest-neighbour (KNN) graphs followed by Leiden community detection. Candidate KNN graphs were constructed across combinations of distance metric (Euclidean or cosine), retained principal components (10, 20, 30 or 50), and neighbourhood size *K* (30, 50 or 80; Table S3). Neighbour distances were converted into edge weights using a radial-basis function,

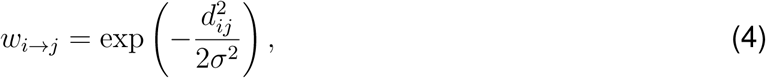

where *d_ij_* denotes the distance between analyses *i* and *j*, and *σ* is the 90th percentile of neighbour distances for the candidate graph. Directed edges were combined into an undirected graph, retaining the larger weight when both directions were present.

To ensure that graphs were well connected and did not group samples by assay technical variants, candidate graphs were required to meet the connectivity thresholds (at least 99% of analyses in the largest connected component and at most two components). We then measured how often profiles were linked to others from the same platform family beyond what was expected by chance, and retained graphs within 0.05 of the lowest value in that root (see Quantification and statistical analysis). Within this eligible set, graphs were ranked by configured preferences for cosine distance, larger neighbourhood sizes, and 30 or 50 principal components, followed by platform mixing (Table S3). Morphology neighbourhood preservation was recorded as a diagnostic measure. Up to six candidate graphs per root were carried forward.

The retained candidate graphs were evaluated by Leiden clustering using the constant Potts model (CPM) objective^52^,

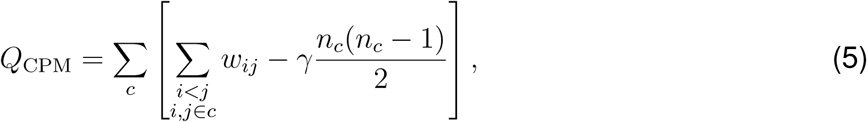

where *c* indexes communities, *w_ij_* is the undirected edge weight between analyses *i* and *j*, *n_c_* is the number of analyses assigned to community *c*, and *γ* is the Leiden resolution parameter. For each graph, *γ* was scanned across a two-stage logarithmic grid using five random seeds per resolution (Table S3). Candidate partitions were evaluated for between-seed reproducibility, community-size balance, low platform association, and stability across adjacent resolutions as defined under Quantification and statistical analysis.

Up to six graph–resolution candidates per root were then evaluated across 20 repeats of 80% subsampling within the fixed principal-component space. When a root had candidates from a stable resolution plateau that passed all subsampling criteria, the final setting was selected from that plateau (three of the 19 released roots). When no plateau candidate passed every threshold simultaneously, the highest-ranking candidate from the subsampling evaluation was retained as the best-available setting (16 roots; ranking rules in Table S3). Selecting the root-level clustering parameters and admitting individual communities as reference subtypes were treated as separate steps.

Each root partition was next refined by evaluating community pairs that were close in the comparison space and repeatedly clustered together across subsampling runs. Such pairs became merge candidates when either community was unstable, when their median genome-wide CNV fractions differed by at least 0.10, or when either community was technically dominated (at least 80% of its analyses from one platform or source, with at least 1.5-fold enrichment). Candidate merges were applied only when they did not cross boundaries supported by reproducible chromosome-scale or gene-level signature differences (Table S3). This produced 142 refined communities across the 20 evaluated roots (139 across the 19 retained atlas roots and three in C47).

Refined communities entered the final atlas through separate core and conditional admission procedures (Table S3). Core admission required at least 200 defining analyses, recovery in at least 90% of subsampling repeats, and the community recovery thresholds. None of the three communities in C47 met these criteria, leaving 103 core subtypes across 19 roots.

Communities recovered in fewer than 90% of subsampling repeats were evaluated for the conditional layer when they showed consistent membership whenever detected and carried both chromosome- and gene-level signatures before release filtering. These communities served as secondary references for profiles that could not be confidently assigned to a core subtype. To verify their stability, each candidate was tested across ten random 80/20 splits, using 80% of its analyses as the reference and projecting the held-out 20% after core evaluation. Admission required held-out members to be consistently and specifically reassigned to their own candidate (≥ 50% coverage and ≥ 90% agreement and precision) while accepting at most 5% of core-subtype controls (Table S3); all nine released conditional subtypes passed all ten repeats.

#### CNV signatures, segment spans and subtype annotations

Each reference subtype was characterized by chromosome-scale (Track 1) and gene-level (Track 2) copy-number signatures relative to other eligible analyses in the same anatomical root, excluding sex chromosomes.

Track 1 summarized chromosome-scale gain and loss patterns directly from the autosomal CNV segments. Within each analysis *i* and autosome *a*, overlapping segments of direction *e* ∈ {gain, loss} were merged, and the covered chromosome fraction was calculated as

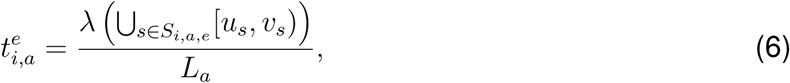

where *S_i,a,e_* is the set of segments of direction *e* on chromosome *a* in analysis *i*, *u_s_* and *v_s_* are segment boundaries, *λ* denotes union length, and *L_a_* is the GRCh38 chromosome length. These fractions measure the chromosomal span of reported CNV segments relative to total chromosome length. This yielded 44 candidate features per analysis (gain and loss across 22 autosomes), with each subtype summarized by the median coverage fraction across its defining analyses.

Track 2 characterized gene-level CNV patterns across the 1,263 autosomal cancer-associated genes in four channels: gain, high-level gain, loss and high-level loss. To describe each gene relative to the sample’s average CNV coverage in the same channel, the mean across the gene set *G* was subtracted from each gene’s coverage fraction *f^k^_ig_*:

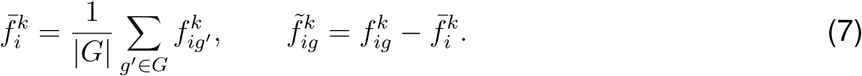

Statistical selection and cross-platform consistency criteria for both tracks are specified under Quantification and statistical analysis (Table S3). Only signatures enriched in a subtype on the comparison scale of the corresponding track (higher chromosome coverage for Track 1; higher mean-centred gene values for Track 2, which does not require a higher unadjusted event frequency) were included in the final release, whereas depleted contrasts were excluded.

Enriched Track 2 events were linked back to the autosomal CNV segments to characterize the genomic span of their supporting alterations. Gain and loss events were matched to same-direction segments, including high-level segments, whereas high-level events were matched only to same-direction high-level segments. An analysis was counted as a segment carrier when matching segments jointly covered at least 50% of the gene interval. Events with no segment carriers were excluded from the release. For each carrier analysis, the shortest overlapping matched segment, expressed as a fraction of chromosome length, represented its span. Supporting segments were classified as focal (at most 2% of the chromosome), regional (more than 2% and less than 50%), or broad (at least 50%). At the subtype level, a released event was classified as mixed when at least two span categories each accounted for at least 25% of carrier analyses, and otherwise took the category with the largest carrier fraction. Events with segment carriers in fewer than 5% of subtype analyses kept their span class and were flagged as low support.

To evaluate whether mixed classifications reflected assay differences, mixed Track 2 events were re-examined within platform families and individual assay platforms having at least ten segment carriers (Table S3). Events represented in at least two such groups were assessed. Within each evaluated platform group, the 25% mixed rule was applied first; otherwise, assigning a focal, regional or broad label required at least 60% of carriers in that category, with remaining groups labelled unresolved. This platform-stratified check assessed the persistence of mixed spans within platforms and differences in carrier spans between platforms.

Each subtype was also annotated with its ICD-O topography and morphology, NCIt diagnosis, assay-platform, and data-source composition. Diagnostic and data-source annotations were descriptive and did not influence signature selection, whereas platform-family annotations were used in the cross-platform support filter described below.

#### Calibrated projection and reference queries

The released projection reference preserved the CNV features, root-specific preprocessing transformations, reference analysis coordinates, subtype identities and subtype-specific calibration limits. Query profiles were encoded using the same 1,263 autosomal genes and transformed into the comparison space of their known anatomical root. Projection used the root’s selected distance metric, number of principal components and *K*. From the *K* nearest reference profiles, we calculated local support (the fraction belonging to the leading candidate subtype), vote margin (the difference between the two highest subtype vote fractions) and the mean distance to neighbours supporting the leading candidate subtype.

Projection limits were calibrated separately for each subtype by leave-one-out projection of its definition analyses, using only analyses correctly reassigned to their original subtype. Calibration required at least 20 correctly recovered analyses and leave-one-out label agreement of at least 80% (Table S3). Among correctly recovered analyses, the 10th percentiles of support and margin set the minimum limits, and the 95th percentile of distance set the maximum accepted distance, *d*^max^_*c*_. A query received an assignment only when all three limits were met. Queries within the distance limit but failing the support or margin requirement were reported as ambiguous; queries exceeding the distance limit or lacking eligible calibration were unresolved. Queries were evaluated first against the core layer; ambiguous or unresolved queries were then evaluated against the conditional layer when one was available.

The application workflow accepts GRCh38 autosomal CNV segments with sample identifiers and anatomical-root metadata. Assigned queries can retrieve subtype diagnostic composition, Track 1 and Track 2 annotations, available mutation-context summaries, nearest reference profiles, and candidate cell-line models (Figure S1). Mutation summaries describe the reference subtype within a diagnosis. If an OncoTree code is supplied, mutation-context rows are filtered to that diagnosis; otherwise, all available diagnosis-specific rows are returned, and no diagnosis is inferred. Records of the same analysis or individual as the query are excluded from retrieved subtype summaries. Cell-line queries exclude their own Cellosaurus accession from the retrieved candidate models.

#### Reconstruction and held-out projection

Reduced-data reconstruction assessed recovery of the 103 core subtypes. Analyses in each root’s fixed principal-component space were divided into five folds balanced by original community assignment. Each fold was omitted once, and the KNN graph and Leiden communities were rebuilt on the remaining analyses using the full-atlas coordinates and selected parameters. The minimum community-size threshold was scaled to the retained fraction; preprocessing and principal component analysis (PCA) were not refitted. Rebuilt communities were mapped to full-data refined communities according to the largest number of shared analyses. Both signature tracks were then recalculated using the original extraction rules. Subtype, membership and signature recovery are defined under Quantification and statistical analysis.

Held-out projection performance was evaluated using one held-out fold (30,609 queries) from a five-fold split of core definition analysis records, balanced within each subtype by platform family and data source. Preprocessing transformations were refitted after excluding the heldout queries. The projection reference and assignment limits were then constructed from the remaining 122,424 core defining analyses, retaining the released subtype labels and selected root-specific parameters. Held-out queries did not contribute to fitting the transformations or calibration limits. Because the split was made at the analysis-record level, 4,986 queries (16.3%) shared an individual identifier with a training reference analysis in the same root. When these queries were removed from the evaluation, without refitting, 21,887 of the remaining 25,623 queries (85.42%) were assigned and 21,877 of these (99.95%) matched their released labels.

To evaluate source-held-out reconstruction, all TCGA source records were excluded before rebuilding the references. The test registry contained one eligible record per participant and anatomical root, selected as the first record by analysis identifier (*n* = 10,156); 323 additional TCGA records were used in neither training nor testing. To prevent patient overlap in training, we also excluded non-TCGA records sharing a TCGA individual or biosample identifier, as well as the GLASS glioma cohort, which includes reprocessed TCGA cases. All remaining non-TCGA records were retained for training. Encoded gene-level features were reused, but the platform-family eligibility screen, preliminary quality control, normalization, standardization and PCA were repeated using the training records. Graph and resolution selection, community refinement, core-reference construction and calibration were repeated with the original search space, with parameters reselected for each root. C47 was predeclared as a root-level group and was not rebuilt, and C49 stopped at the start of community refinement, leaving 193 held-out TCGA records in these roots without a rebuilt subtype reference.

Held-out TCGA records were projected into their corresponding rebuilt anatomical roots. Assignment coverage used the full test registry as the denominator. Grouping correspondence was evaluated for records assigned to a rebuilt subtype that also carried a released core-subtype label (*n* = 6,981).

#### Somatic mutation associations and cross-cohort comparison

We compared somatic mutation frequencies between core CNV subtypes within the same anatomical root and OncoTree diagnosis. Mutation and clinical data came from AACR Project GENIE release 17.0-public^50^. Progenetix CNV analyses were matched to GENIE sample identifiers and linked to the official sample and patient tables.

We selected one defining CNV analysis per patient and anatomical root before examining mutation associations. Primary samples were preferred, followed by younger reported age at sequencing. Lexical sample and analysis identifiers broke remaining ties. Only selected analyses belonging to core subtypes were retained. If the selected analysis belonged to a conditional subtype, it was excluded without selecting a replacement. A gene was evaluable only if the assay declared coding-exon coverage and included an exon of that gene in its GENIE panel-design file.

Mutation calls required FILTER=PASS, Annotation_Status=SUCCESS and NCBI_Build=GRCh37, and one of the nine variant classes listed in Table S3. A qualifying somatic call made the sample mutation-positive for that gene. If qualifying calls were exclusively non-somatic, the gene state was uncertain and the sample was excluded for that gene. With no qualifying call, the sample counted as mutation-negative only if it had at least one raw MAF record. Samples without any raw MAF record were excluded.

Before mutation-status filtering, 43,096 unique core-subtype analyses met the root, diagnosis, gene-panel and subtype requirements for the comparisons. Of these, 2,842 (6.59%) lacked raw MAF records. In total, 40,254 unique analyses were evaluable for at least one comparison, spanning 97 core subtypes and 19 roots. Mutation and local CNV features were matched by gene symbol.

For each gene, the target subtype was compared with the pooled remaining eligible core subtypes in the same root and diagnosis. The target and each control subtype required at least 20 evaluable analyses. This defined a fixed family of 9,582 comparisons. All three association models were estimable for 3,951 comparisons involving 81 core subtypes in 16 roots. Model specifications are given under Quantification and statistical analysis.

We then examined the 45 associations retained across the three GENIE models using public TCGA MC3 v0.2.8 mutation calls^51^. One TCGA SNP6 defining analysis (platform geo:GPL6801) was selected per participant and root. Selection preferred sample type 01, then vial A, followed by lexical barcode and analysis identifiers. Selected conditional-subtype analyses were excluded without replacement. Each Progenetix TCGA record was paired with a unique MC3 tumour aliquot using the exact 16-character sample-vial barcode.

For TCGA, a qualifying PASS call defined mutation-positive status. If qualifying calls appeared only under other filters, the gene state was unknown. With no qualifying call, mutation-negative status required at least one PASS record in MC3 for the matched aliquot. Otherwise, the state was unknown.

Seven OncoTree diagnoses were matched to TCGA projects using morphology, topography and diagnosis-label filters (Table S3): colon adenocarcinoma (COAD), rectal adenocarcinoma (READ), lung adenocarcinoma (LUAD), invasive ductal breast carcinoma (IDC), prostate adenocarcinoma (PRAD), bladder urothelial carcinoma (BLCA) and clear-cell renal cell carcinoma (CCRCC). The broad parent code BRCA was not mapped to a single TCGA cohort. GBM was not tested because its diagnostic-code mapping remained unresolved.

Each target and control subtype required at least 20 paired participants. Only control subtypes represented in both cohorts were used. GENIE estimates were recalculated using this same control set before association directions were compared.

#### Cell-line recommendation and signature representation

A cell line could have several CNV profiles in the catalogue. We projected the 4,605 eligible profiles using the hierarchical core-to-conditional procedure. Profiles were then grouped by curated Cellosaurus accession and anatomical root. Recommendations were made separately for each cell-line–root group.

Within each group, the subtype receiving the most assignments was called the consensus subtype. Assignment coverage was the fraction of all available profiles receiving a calibrated assignment. Assignment consistency was the fraction of assigned profiles supporting the consensus. Among profiles supporting the consensus, the representative had the smallest mean distance to reference neighbours of that subtype.

A model with one available profile was recommended when that profile received a calibrated assignment (Tier B). For models with multiple profiles, at least two had to support the consensus subtype. Recommendation also required at least 50% assignment coverage and 60% assignment consistency (Tier A). Other assigned groups were provisional when consistency exceeded 50%, and discordant otherwise. Profiles were catalogue records and were not treated as independent biological replicates.

Models recommended for the same subtype were ranked by evidence tier, assignment consistency, median relative distance and median vote margin. Higher consistency, lower relative distance and higher vote margin were preferred. For each model, both medians were computed using the profiles that supported the consensus subtype. Relative distance was *d*_relative_ = *d*_query_/*d*^max^_*c*_, where *d*_query_ was the mean distance to reference neighbours supporting the assigned subtype. Relative distances at or below 1 met the subtype-specific distance limit. Figure S1 provides a worked example of retrieval, projection and cell-line queries.

To identify gaps in CNV representation, we checked each subtype signature against the profiles of its matched cell-line models. All available profiles within the corresponding root were included, even when they did not support the model’s consensus assignment.

Track 2 events were recorded when the gene-level CNV fraction exceeded zero in the corresponding direction and amplitude channel; high-level gain and loss events were also counted in the any-amplitude channels. An event was non-evaluable in a profile when source segments, gene rows or finite gene values were missing; high-level events additionally required amplitude-level segment annotations. Non-evaluable profiles were marked as unknown and were not counted as zero.

We then summarized Track 2 evidence across the profiles of each model. Model–event pairs with no evaluable profiles were classified as unknown and excluded from the all/none/some categories. Unknown-profile counts were retained for each pair. Track 1 evidence used continuous chromosome coverage from the union of same-direction segments. For both tracks, continuous values were summarized by the median across evaluable profiles within each model.

Figure 7C summarized signature coverage by recommended models. Each annotation specified a gene, CNV direction and amplitude within one subtype. Annotations were counted separately even when they shared a CNV segment. The analysis considered released Track 2 annotations whose unadjusted event frequency was higher than in the root background. There were 2,942 such annotations; 2,210 belonged to subtypes with recommended models and formed the evaluated set (Table S4).

An annotation was covered if the event was recorded in at least one evaluable profile from any recommended model. The remaining annotations fell into two groups: those recorded only in other matched models, and those unrecorded in every matched model of that subtype. For the latter group, we distinguished annotations with incomplete evaluability from those with fully evaluable profiles but no recorded event. Different recommended models could contribute coverage for different annotations.

In Figure 7B, reference gain and loss values were mean encoded scores across defining analyses, divided by three. Scaled values below 0.05 were displayed as zero. Cell-line profiles used the representative recommended profile. Figure 7D compared chromosome-loss coverage in each model profile with the median across defining tumour analyses. Recording of *BIRC3* loss was shown separately. The QGP-1 comparison used its single available catalogue profile.

### Quantification and statistical analysis

In CNVSIM, graph and community metrics informed reference construction. Signature tests and mutation models supplied subtype annotations, while reconstruction and projection metrics evaluated the selected reference. Selection and support thresholds are listed in Table S3.

Unless otherwise stated, *n* is the number of Progenetix CNV analysis records. Mutation comparisons used one selected record per patient or participant and anatomical root. Cell-line recommendations used Cellosaurus-accession–root groups.

#### Graph, community and conditional-reference evaluation

Graph diagnostics screened candidate KNN graphs before Leiden clustering (Figure 2A). Connectivity was measured by the number of components and the fraction of records in the largest component. Platform clustering was measured as the excess fraction of same-platform-family neighbours above its random-mixing expectation. For *n* records and *K* neighbours per record,

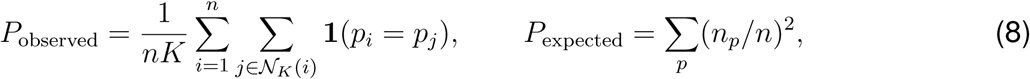

where *p_i_* is the platform family of record *i*, and *n_p_* is the number of records in that family. Their difference was Δ_platform_ = *P*_observed_ − *P*_expected_. Morphology preservation was the fraction of neighbours sharing the recorded morphology label.

Leiden partitions were then evaluated for reproducibility and community-size balance. Seed stability was the mean pairwise adjusted Rand index (ARI) across partitions from different random seeds^53^. Small-community mass was the fraction of records in communities below the minimum size in Table S3. The largest-community fraction was max*_c_ n_c_/n*, where *n_c_* is community size. The representative partition minimized small-community mass, with CPM quality as the tie-breaker.

Platform association was also assessed at the community level. Bias-corrected Cramér’s *V* quantified association between community assignment and platform family^54^. It used SciPy’s chi-squared statistic from the community-by-platform table, with continuity correction for 2 × 2 tables. Lower values indicated weaker association. Stable plateaus required agreement between partitions at adjacent eligible resolutions; eligibility, agreement and plateau-length thresholds are listed in Table S3.

Subsampling assessed whether candidate partitions persisted after records were omitted. Evaluation coverage was the fraction of retained analyses assigned to non-outlier communities in both the full-data and reconstructed partitions. Candidate selection used the mean coverage across repeats. Whole-partition agreement used ARI and the fraction matched by one-to-one community matching that maximized total shared membership.

To assess member retention and mixing, each full-data community was separately matched to the rebuilt community with the greatest overlap. Let *C* and *M* denote their evaluated member sets, with *O* = |*C* ∩ *M* |. Then

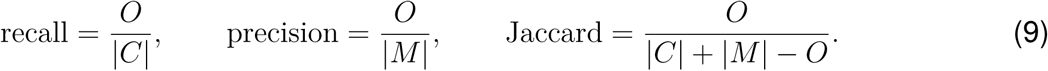

For recall, the tenth percentile across communities was calculated within each repeat. Candidate selection used the median of these values across repeats. For precision and Jaccard similarity, each community’s tenth percentile across repeats was calculated, and the lowest community value was used (Table S3).

For refined-community admission, rebuilt communities mapping to the same refined community were pooled. Detection was the proportion of repeats in which a community was recovered; recovery metrics were summarized across detected repeats.

For conditional admission, held-out candidate queries were evaluated after an ambiguous or unresolved core assignment. Coverage was the fraction receiving any conditional assignment. Label agreement was the fraction of assigned eligible queries returning to their original candidate. Precision was the fraction of non-core queries accepted by a candidate that were its held-out members. Its denominator included accepted queries from other conditional candidates and other non-core analyses routed to conditional evaluation within the same root.

Acceptance of core-subtype profiles was checked separately by projecting sampled same-root core controls directly against the conditional reference. For each candidate, core-control acceptance was the fraction of these controls assigned to it.

#### Signature statistics and platform sensitivity

Signature tests selected chromosome-scale and gene-level subtype annotations (Figure 3). They also supplied provisional signature evidence for community refinement. Two-sided Mann– Whitney *U* -tests compared each subtype with its same-root background, using chromosome coverage for Track 1 and within-analysis mean-centred gene fractions for Track 2. The rank-based effect was AUC = *U/*(*n_c_n_,¬,c_*), with subtype records as the first group and background records as the second. AUC is the probability of a higher subtype value, with ties counting one-half. Table S3 lists the effect-size, candidate-count and platform-support thresholds. Platform eligibility required both subtype and background groups to meet the sample-size minimum.

For Track 1, Benjamini–Hochberg correction covered the 44 chromosome-direction features separately for each subtype. The pooled effect was 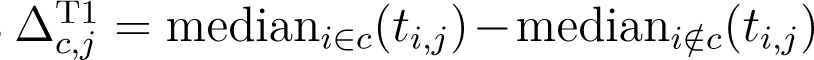. Selection required *q* ≤ 0.05 and the specified minima for the absolute median difference and absolute AUC shift from 0.5. At least two eligible families had to show same-direction median differences meeting the family-specific minimum.

For Track 2, a gene-channel comparison was excluded if both groups had fewer than max[3, ⌈0.01 min(*n_c_, n_,¬,c_*)⌉] non-zero original values. At least two valid adjusted values were required per group. The same rule was applied within eligible platform families using their group sizes. Benjamini–Hochberg correction covered tested genes separately for each subtype and channel. Adjusted mean differences and unadjusted frequency differences were reported separately:

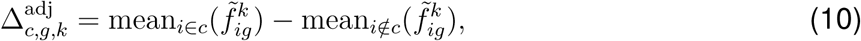

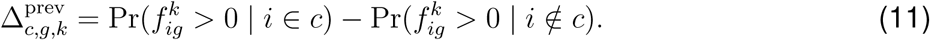

Candidates required *q* ≤ 0.05 and either the absolute frequency-difference or absolute AUC-shift minimum. AUC shifts were measured from 0.5. Positive adjusted effects additionally required AUC at least 0.60; negative effects required AUC at most 0.40. Within each subtype, channel and direction, candidates were ranked by adjusted *P* -value, unadjusted frequency difference and AUC. The specified candidate cap was applied before platform evaluation.

Cross-platform evaluation assessed whether pooled gene-level signals were also supported within individual platform families. Within a family, support required the adjusted mean difference and AUC to agree with the pooled direction. Either the absolute frequency difference or the absolute AUC shift also had to meet its minimum (Table S3). Events were selected with support from at least two families and at least two-thirds of evaluated families. Across all evaluated families, non-zero adjusted effects had to agree in direction and the range of AUC values had to meet the specified limit.

For the mixed-span platform check described above, Kruskal–Wallis tests assessed differences in carrier-segment spans across eligible platform groups. Epsilon-squared quantified their magnitude as max[0, (*H* − *k* + 1)*/*(*N* − *k*)], where *H* is the test statistic, *k* the group count and *N* the carrier count. Benjamini–Hochberg correction covered tested events separately for platform-family and individual-platform comparisons. Platform-associated differences required *q* ≤ 0.05 and the minimum effect in Table S3.

#### Mutation association models and sensitivity analyses

We asked whether subtype–mutation associations persisted after accounting for sequencing assay and CNV at two scales: overall genomic burden and copy-number status at the tested gene (Figure 4; Table S5). Overall burden was represented by FGA, the reported fraction of the genome affected by CNV. Subtypes differ in overall CNV burden, and burden may itself relate to mutation frequency, so an unadjusted subtype difference could reflect burden. We therefore used three main models to evaluate the same subtype contrast under different representations of these global and local CNV features.

The primary model compared records from the same assay with similar overall CNV burden and the same binary gain/loss state at the tested gene. In GENIE, FGA values were ranked within each diagnosis–gene cohort and divided into five groups. These FGA quintiles, sequencing assay and the joint binary local gain/loss state defined the strata. Local gain and loss were represented by 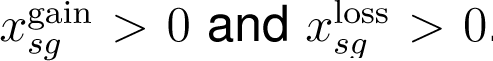. A stratum was informative when it contained target and control analyses and both mutation-positive and mutation-negative outcomes.

Within each informative stratum, the exact test held the target/control group sizes and total mutation-positive count fixed. The resulting hypergeometric distributions were combined by convolution to obtain the null distribution of the total positive target count. The two-sided exact *P* -value summed probabilities at least as far from the null expectation as the observed total. Association direction, magnitude and uncertainty were summarized by Mantel–Haenszel common odds ratios and 95% confidence intervals from statsmodels.StratifiedTable.

The second main model tested whether subtype–mutation associations remained when high-level gains and losses were distinguished from other gains and losses. This amplitude model used paired encoded scores (0, 1 or 3), while retaining the assay and FGA-quintile stratification. Both stratified models required the informative-stratum and target/control support criteria in Table S3, including a finite exact *P* -value.

The third main model tested whether the subtype–mutation association remained after replacing FGA groups with continuous burden adjustment. The continuous-FGA model used binomial logistic regression with a restricted cubic spline for FGA and fixed effects for each assay-by-binary-local-CNV combination, using records from the primary model’s informative strata. Spline knots were calculated from the complete diagnosis–gene cohort in GENIE and the complete comparison cohort in TCGA, using the percentiles listed in Table S3.

In this regression, *T_i_* indicated target-subtype membership and *β* was its coefficient. The adjusted mutation odds ratio between the target subtype and its controls was exp(*β*), with a 95% Wald confidence interval of exp[*β* ± 1.96 SE(*β*)]. A one-degree-of-freedom likelihood-ratio test compared the model with and without *T_i_* to assess whether subtype membership remained associated with mutation status after these adjustments. Logistic fits required at least five mutation-positive and five mutation-negative records and the minimum residual degrees of freedom in Table S3, as well as distinct spline knots, an identifiable subtype effect, convergence and finite stable estimates.

Within GENIE, Benjamini–Hochberg adjustment covered all 9,582 comparisons separately for each main and sensitivity model, with non-estimable comparisons entered as *P* = 1. An association entered the reported set only if all three main models were estimable, all three had *q <* 0.05, and their odds ratios agreed in direction. We then assessed the robustness of these associations to additional mutation and sampling variables, sample exclusions and differences between centres.

Seven additional logistic models started from the continuous-FGA regression described above (Table S3). On the same complete primary/metastatic records, a baseline model provided a comparison for models adding mutation density, sample type or both. Three variants of the combined density-and-sample-type model then used local gain/loss amplitudes, added a quadratic term in transformed density, or restricted the sample set.

Mutation density counted qualifying somatic calls outside the tested gene per megabase of the assay’s merged exonic target regions and entered as log(1 + density). Models used informative strata identified on complete primary/metastatic records under the primary stratification. The variant using gain/loss amplitudes also had to meet the data-support criteria of the stratified amplitude main model (Table S3).

The restricted variant examined whether associations persisted after excluding samples with high mutation density or qualifying somatic calls in mismatch-repair or polymerase genes other than the tested gene (Table S3). Informative strata were reassessed after these exclusions.

For the reported associations, retention after additional adjustment was evaluated using the model that included both mutation density and sample type. Retention required *q <* 0.05 and an odds ratio in the same direction as the primary model.

A separate centre analysis examined whether associations differed between MSK and non-MSK records. Models were fitted within each group, and a centre-by-subtype interaction likelihood-ratio test assessed differences between their subtype effects. The interaction test required both groups to meet primary-model support; its *P* -values were Benjamini–Hochberg-adjusted across the 45 reported associations.

Finally, the three main models were applied in TCGA to compare association directions across cohorts. Project replaced sequencing assay, and FGA quintiles were formed within each comparison cohort, with ties broken by analysis identifier. Benjamini–Hochberg adjustment used the fixed 45-candidate family separately for each model, with unavailable or non-estimable comparisons entered as *P* = 1. Direction agreement was reported separately from significance at *q <* 0.05 across all three TCGA models.

#### Reconstruction and projection metrics

Reduced-data reconstruction was evaluated at subtype, membership and signature levels (Figure 6A). A released core subtype was recovered when at least one rebuilt community mapped to it by membership overlap. Membership recovery was the proportion of retained core defining analyses whose rebuilt community mapped to their released subtype. Exact signature recovery required the same chromosome, CNV direction and enrichment direction for Track 1. For Track 2, it required the same gene, CNV channel and enrichment direction.

Exact signature recovery measured which events were selected again. Enrichment-profile similarity assessed whether their relative effect patterns were preserved after rebuilding. For each subtype and track, we formed a complete-atlas vector **u** and a rebuild vector **v**, with matching entries for the released events. Entries were 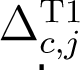 for Track 1 and 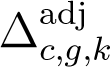 for Track 2. Rebuild values came from the complete fold statistics, including events that were not selected again. Estimates unavailable because of an unrecovered subtype or insufficient observations were set to zero in the rebuild vector.

We compared these vectors using cosine similarity, **u**^T^**v***/*(∥**u**∥_2_∥**v**∥_2_). Higher values indicated closer agreement in the relative effect pattern. Similarity was calculated when at least two released events were available and the rebuild vector was non-zero. Subtype-level values were summarized by the median within each root.

We then evaluated query assignment in the internal held-out projection test. Assignment coverage measured the fraction of queries receiving an assignment, *N*_assigned_*/N*_queries_. Label agreement was the fraction of assigned queries matching their released subtype label. Unassigned queries contributed to the coverage denominator.

TCGA source-held-out reconstruction could change subtype boundaries and identifiers. We therefore compared how pairs of records were grouped. The comparison set comprised records with both a released core-subtype label and a rebuilt subtype assignment. Within each root, *n_ab_* counted records with released label *a* and rebuilt label *b*. Row totals were *n_a,·,_*, column totals were *n_,·,b_*, and the total was *n*. The fraction of same-released-subtype pairs kept together was

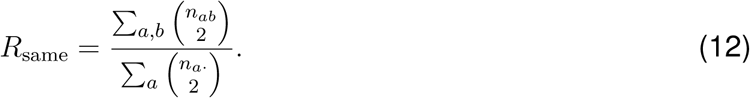

The fraction of different-released-subtype pairs kept apart was

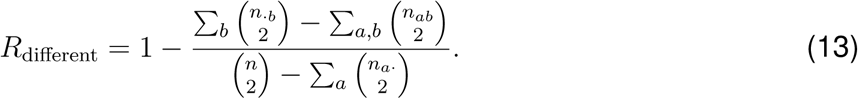

Lower *R*_same_ indicated splitting of released subtypes; lower *R*_different_ indicated merging between them. ARI and adjusted mutual information provided complementary summaries of partition agreement in the validation tables. Figure 6C flags roots with fewer than ten compared records in any represented released subtype; the validation tables also flag comparisons with fewer than 30 records.

## Notes

### Competing Interest Statement

The authors have declared no competing interest.

### Summary of Updates

Manuscript text and title revised for clarity; within-diagnosis somatic mutation association analyses in GENIE and cross-cohort evaluation in TCGA added; TCGA-excluded atlas reconstruction and projection validation added; cancer cell-line evaluation expanded to assess gene-level CNV signature coverage gaps; Figures 1 to 7 reorganized and updated; Methods expanded to detail all statistical models and metrics; Supplemental Tables S1 to S5 and CNVSIM v1.0.1 data and code release updated.

