## Supplemental Figure 1 for "CNVSIM: an interpretable cross-study copy-number reference atlas for cancer subtyping"

### CNVSIM applications

Run from the repository root. $CNVSIM_DATA: complete data release; $EXAMPLE_OUT: new output directory.

#### Inputs

#### Outputs

#### A. Project a CNV sample and retrieve its reference context

##### 1. Find analyses for a public biosample

```
python \
  tests/progenetix_retrieval_example/query_candidates.py \
  --resource tumour --biosample-id pgxbs-kftvhy2d \
  --limit 20 --release-dir "$CNVSIM_DATA" \
  --out "$EXAMPLE_OUT/candidates.tsv"
```

```
[CNVSIM] bycon candidate query complete
  resource: tumour
  CNV analyses: 1
  ready for selection: 1
  metadata unavailable: 0
  Mark selected=yes for every analysis to retrieve and project.
```

Set selected=yes in candidates.tsv for the analysis to use.

candidates.tsv after selection (selected columns)

| selected | retrieval_status | biosample_id | analysis_id |
|---|---|---|---|
| yes | ready | pgxbs-kftvhy2d | pgxcs-kftw1ui5 |

##### 2. Download CNV segments from Progenetix and prepare inputs

```
python \
  tests/progenetix_retrieval_example/prepare_projection_inputs.py \
  --selection "$EXAMPLE_OUT/candidates.tsv" \
  --out-dir "$EXAMPLE_OUT/input"
```

```
[CNVSIM] projection inputs prepared
  analyses: 1
  autosomal CNV segments: 187
```

Creates samples.tsv and segments.tsv in input/.

input/samples.tsv (all columns); segments.tsv: 187 rows

| sample_id | icdo_topography_id | genome_build |
|---|---|---|
| pgxcs-kftw1ui5 | pgx:icdot-C50.9 | GRCh38 |

##### 3. Project the sample and retrieve context

```
python run_scripts/applications/query_reference_context.py \
  --release-dir "$CNVSIM_DATA" \
  --samples "$EXAMPLE_OUT/input/samples.tsv" \
  --segments "$EXAMPLE_OUT/input/segments.tsv" \
  --out-dir "$EXAMPLE_OUT/context"
```

```
pgxcs-kftw1ui5: projected_assignment | C50-CTX0 | query_001_pgxcs-kftw1ui5/report.json
```

Subtype assignment and cell-line summary
(from projection_assignments.tsv & report.json)

| Assigned subtype | Supporting neighbours | Mean distance | Matched cell lines | Recommended cell lines |
|---|---|---|---|---|
| C50-CTX0 | 62/80 | 0.3071 | 14 | 9 |

14 Cellosaurus cell lines match subtype C50-CTX0; 9 meet CNV recommendation criteria.

#### Cell-line CNV evidence and tumour mutation context for assigned subtype C50-CTX0 (from Step 3)

##### 1) How is MYC gain represented across the 9 recommended cell lines?

| Recommended models | Recorded in all evaluable profiles (recorded_only) | Not recorded in any evaluable profile (not_recorded_only) | Profiles disagree (mixed) | No usable profiles (unknown_only) |
|---|---|---|---|---|
| 9 | 6 | 2 | 1 | 0 |

BT-20 has 8 CNV records: 4 record MYC gain, 2 do not, and 2 lack CNV data.

| Cell line | Profiles with MYC gain | No MYC gain recorded | CNV data unavailable | Evidence status |
|---|---|---|---|---|
| HCC1937 | 18 | 0 | 0 | recorded_only |
| CAL-85-1 | 0 | 3 | 0 | not_recorded_only |
| BT-20 | 4 | 2 | 2 | mixed |

##### 2) Which somatic gene mutations are associated with C50-CTX0 in tumour cohorts?

| Diagnosis | Gene | GENIE effect | GENIE OR | TCGA OR | TCGA status |
|---|---|---|---|---|---|
| BRCA | GATA3 | depleted | 0.1479 | — | cohort_unavailable |
| BRCA | PIK3CA | depleted | 0.3643 | — | cohort_unavailable |
| BRCA | TP53 | enriched | 9.463 | — | cohort_unavailable |
| IDC | ESR1 | depleted | 0.06912 | — | not_estimable |
| IDC | GATA3 | depleted | 0.1335 | 0.09599 | estimable |
| IDC | PIK3CA | depleted | 0.4487 | 0.3897 | estimable |
| IDC | TP53 | enriched | 15.28 | 7.183 | estimable |

These are mutation patterns in groups of tumours, not mutation results for this sample or the cell lines. Both BRCA and IDC are shown because no diagnosis was supplied.

#### B. Query Cellosaurus cell lines (direct catalogue lookup)

##### a) Find recommended cell lines for a known subtype

```
python run_scripts/applications/query_experimental_models.py \
  --cell-line-dir "$CNVSIM_DATA/cell_line" \
  --reference-key C16-COND4 --top-n 5 \
  --out "$EXAMPLE_OUT/C16-COND4_models.tsv"
```

```
CNVSIM cell-line recommendations
Reference: C16-COND4
Layer: conditional | Root: C16 | Models returned: 1
 rank  model accession  tier  profiles consistency coverage distance/limit
   1 GIST522 CVCL_7042 Tier B       1     100.0%  100.0%         0.162
Lower distance/limit indicates a closer calibrated match.
```

##### b) Look up the reference subtype for a known cell line

```
python run_scripts/applications/query_experimental_models.py \
  --cell-line-dir "$CNVSIM_DATA/cell_line" \
  --model-id CVCL_7042 --include-analysis-details
```

```
CNVSIM cell-line lookup
Model: GIST522 (CVCL_7042)
Root: C16
Match: C16-COND4 [conditional] | Status: projected_consensus
Evidence: Tier B | Single-profile assignment
Profiles: 1 | assigned: 1 | consensus support: 1
Consistency: 100.0% | coverage: 100.0% | distance/limit: 0.162
```
